# Topology-aware reconstruction of cellular state landscapes from microscopy using self-supervised learning

**DOI:** 10.64898/2026.05.30.728966

**Authors:** Elisa Messori, Doaa Taha, Lisa Fournier, Anna Foix Romero, Virginie Uhlmann, Pascal Frossard, Cédric Vincent-Cuaz, Rickie Patani, Raphaëlle Luisier

## Abstract

Morphology and spatial organisation provide complementary readouts of cellular state. However, reconstructing continuous cellular state landscapes from imaging data remains challenging, particularly in dense biological cultures. Here we present SI-SimCLR, a spatially informed self-supervised learning framework that learns biologically informative representations directly from fluorescence microscopy images without requiring segmentation or manual annotation. Combined with a graph-based partial optimal transport framework, SI-SimCLR enables reconstruction of cellular phenotypic landscapes from static imaging data, revealing how phenotypic substates are organised and connected. To establish and validate this framework, we generated a multimodal dataset of human iPSC-derived astrocytes using high-content imaging and matched bulk transcriptomics. SI-SimCLR resolved distinct interconnected astrocyte substates associated with disease and inflammatory states. ALS astrocytes occupied constrained regions of the morphological landscape. Strikingly, morphology and transcriptomics captured distinct and complementary aspects of astrocyte state variation.Together, our framework establishes a scalable and annotation-free strategy for reconstructing cellular phenotypic landscapes from microscopy data, enabling analysis of cellular heterogeneity, landscape connectivity and phenotypic responses across biological systems.

## INTRODUCTION

The rapid expansion of single-cell transcriptomic profiling has enabled the construction of increasingly comprehensive atlases of cellular states, transforming our understanding of development, disease progression and therapeutic response ^1–4^. Yet cellular identity is not encoded solely in molecular programmes. Cell morphology and arrangements capture a rich and complementary layer of phenotypic organisation^5^, integrating cytoskeletal architecture, spatial arrangement, cell–cell interactions and environmental context. Advances in high-content fluorescence microscopy now enable quantitative imaging of millions of cells at high spatial resolution and at substantially lower cost than sequencing-based assays^6^, while providing scalable and minimally disruptive phenotypic readouts compatible with longitudinal disease modelling and high-throughput screening. Together, these developments motivate the generation of new computational frameworks capable of reconstructing cellular state landscapes directly from imaging data.

Recent studies have demonstrated the promise of large-scale imaging-based profiling, including genome-wide perturbation atlases in cultured human cells ^7,8^ and spatially resolved atlases of tumour ecosystems in histopathology ^9^. However, existing approaches for reconstructing cellular state landscapes from images remain fundamentally limited. Most rely on accurate single-cell segmentation, which becomes unreliable in dense or confluent cultures where cell boundaries overlap extensively and morphology itself is highly heterogeneous^10,11^. In parallel, self-supervised learning (SSL) has emerged as a promising alternative to segmentation-based feature extraction in microscopy imaging^12–15^. Yet standard SSL frameworks were primarily developed for visually diverse natural images and often perform suboptimally on biological microscopy data, where samples often exhibit high visual similarity, increasing the risk of focusing on trivial features rather than biologically meaningful variation^16–18^. Furthermore, most existing approaches remain inherently single-cell-centric, neglecting collective spatial organisation and multicellular structural context despite these being integral components of phenotypic identity in many biological systems ^19^.

Beyond cataloguing discrete states, a central objective in developmental and disease biology is to understand how cells are organised within continuous phenotypic landscapes and how cellular states are connected through accessible paths of phenotypic change, as conceptualised by the Waddington landscape paradigm^20^. In transcriptomics, these relationships have been reconstructed using manifold learning, RNA velocity and optimal transport (OT)-based approaches^21–25^. OT methods have proven particularly powerful for characterising the geometry and connectivity of cellular state spaces, enabling reconstruction of developmental trajectories^26^, mapping of cell states across space and time^27^, and prediction of perturbation-induced state transitions^21,25,28^. By contrast, comparable approaches based on cell morphologies and arrangements remain in their early stages, with recent work beginning to quantify transition dynamics from imaging data during directed transdifferentiation and reprogramming^29^. However, these approaches remain limited to constrained biological processes and do not address the more general problem of reconstructing global phenotypic landscapes and accessibility relationships directly from static imaging data. As a result, most imaging frameworks rely on predefined experimental labels, rather than recovering continuous morphological state spaces with emergent attractor-like, intermediary and highly plastic configurations. Reconstructing such phenotypic landscapes from microscopy data requires methods capable of learning representations from dense microscopy data, identifying morphological substates without prior annotation, and inferring their relative accessibility within a unified state space.

These requirements are particularly relevant for astrocytes, whose morphology is tightly coupled to functional state and undergo profound remodelling during neuroinflammation and neurodegeneration ^30–32^. Astrocytes are increasingly implicated across a broad spectrum of neurological disorders, from neurodegeneration^33,34^ to psychiatric disease^35,36^, suggesting that astrocyte state dysregulation may represent a common axis of central nervous system vulnerability^37,38^ with substantial therapeutic potential ^39^. Unlike many cultured cell types, astrocytes display striking morphological heterogeneity, ranging from highly spread to elaborate filamentous configurations, while growing predominantly in confluent arrangements, which further complicate segmentation-based analyses. Importantly, reactive astrocyte states form a continuous and heterogeneous phenotypic spectrum rather than discrete categories, making substate-level resolution essential for understanding how astrocyte diversity contributes to disease. Although transcriptomic studies have begun to resolve astrocyte substates in neurological disorders^33,40–42^, the corresponding morphological landscape remains poorly characterised^43,44^. Consequently, it remains unclear whether morphology recapitulates transcriptionally defined astrocyte states or instead captures complementary aspects of disease-associated heterogeneity.

Here we present SI-SimCLR, a spatially informed self-supervised learning framework for segmentation-free analysis of dense microscopy images. SI-SimCLR extends contrastive learning by incorporating spatially neighbouring image crops as additional positive pairs during training, increasing visual diversity while preserving shared local biological context^17,45,46^. Combined with a graph-based partial optimal transport framework, SI-SimCLR enables topology-aware reconstruction of cellular state landscapes directly from static fluorescence microscopy data without requiring annotation or complementary molecular measurements. Benchmarking against standard SimCLR trained on the same microscopy dataset, as well as supervised and self-supervised models pretrained on natural or fluorescence microscopy images, demonstrated that SI-SimCLR more effectively captures biologically meaningful morphological variation across experimental batches. To establish and validate this framework, we generated a multimodal dataset of human iPSC-derived astrocytes carrying ALS-associated VCP mutations, profiled under basal and inflammatory conditions using high-content imaging and matched transcriptomics. Application of SI-SimCLR revealed that ALS astrocytes occupy constrained and genotype-associated regions of the morphological landscape that partially redistribute toward control-associated configurations following inflammatory stimulation. Strikingly, comparison with matched transcriptomic profiles showed that morphology and gene expression captured distinct but complementary aspects of astrocyte state organisation: inflammatory signalling dominated global transcriptional variation, whereas morphological states remained primarily structured by ALS-associated genotype. These findings indicate that imaging-derived phenotypes capture disease-associated organisation that is only partially reflected by transcriptomic programmes.

Together, our work establishes a scalable framework for reconstructing cellular phenotypic landscapes directly from microscopy images without segmentation or manual annotation. More broadly, it demonstrates how combining spatially informed self-supervised learning with optimal transport-based topology inference enables substate-resolved analysis of cellular heterogeneity across biological systems.

## RESULTS

### Spatially informed contrastive learning for morphological characterisation of microscopy images

To establish and validate a framework for reconstructing continuous cellular phenotypic landscapes from dense microscopy data, we generated a multimodal imaging and transcriptomic dataset of human iPSC-derived astrocytes, a cellular system characterised by pronounced morphological heterogeneity, extensive cell overlap and strong context-dependent remodelling. Astrocytes were derived from control and ALS-associated VCP-mutant lines^47^, cultured under basal conditions or exposed to low- or high-dose pro-inflammatory stimulation (IL-1α, TNF and C1q; TIC) (**Fig. 1A**). Cells were generated across four independent experimental batches spanning distinct culture preparations and imaging sessions.

**Figure 1.**
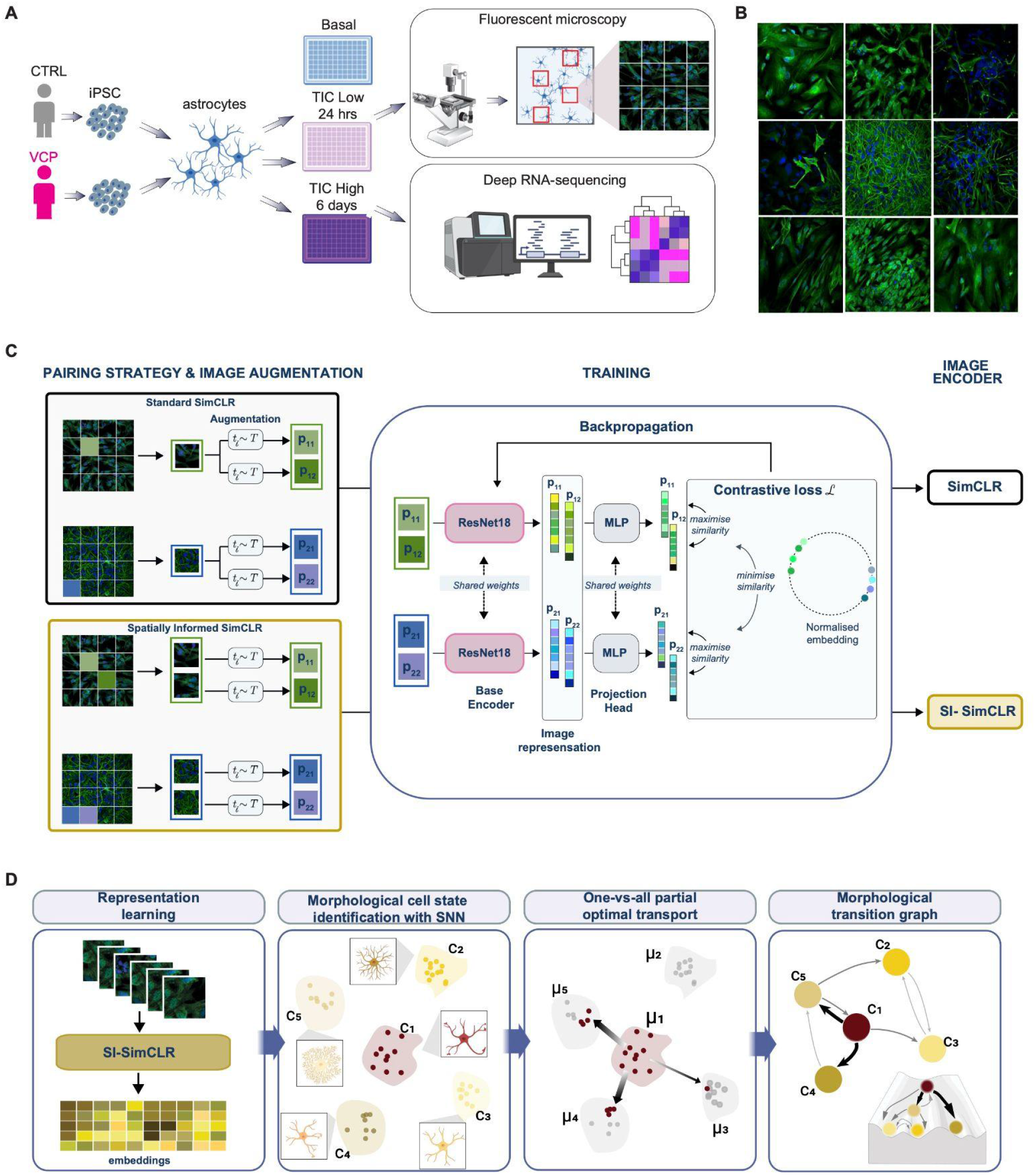
Integrated cellular imaging and transcriptomic atlas of astrocyte phenotypes with contrastive learning-based representation of morphology. **A.** Experimental workflow integrating high-content cellular imaging and deep transcriptomic profiling. Patient-specific hiPSC-derived astrocytes were cultured under multiple conditions to capture a broad spectrum of disease- and inflammation-associated phenotypes at both the cellular and molecular levels. (*Upper*) High-content fluorescence microscopy (DAPI, GFAP) was used to quantify rich cellular features, including morphology, texture, and spatial organisation. Each field of view (red rectangle) was systematically subdivided into sixteen image crops (inset) to enable scalable, fine-grained phenotypic analysis. (*Lower*) Deep bulk RNA sequencing (poly(A)-selected; ≥30M paired-end reads per sample) was performed across 18 samples (see **Supplementary Table 3**), providing high-resolution transcriptomic profiles to comprehensively capture molecular states across conditions. **B**. Representative fluorescence images highlighting the diversity of cellular phenotypes across astrocyte cultures. DAPI and GFAP signals are shown in the blue and green channels, respectively, revealing variation in nuclear organisation, astrocytic morphology, and structural features across conditions. **C**. SimCLR-based contrastive learning framework used to train a ResNet18 encoder for fluorescence microscopy images. In the standard strategy (*black*), a positive pair consists of two augmented views of the same crop. In the Spatially Informed SimCLR (SI-SimCLR; *gold*), positive pairs are formed from adjacent image crops, each independently augmented, introducing contextual variation. **D**. Overview of pipeline for generation of morphological transition graph. Following representation learning with SI-SimCLR, substates are identified via unsupervised clustering. A partial optimal transport (OT) framework is then applied to model the dynamics between the resulting substates. OT-derived costs are used to construct a probability transition matrix to infer plausible transition dynamics between substates.

High-resolution fluorescence imaging combined with cytoskeletal markers such as GFAP has emerged as a powerful approach for quantifying reactive morphological remodelling in astrocytes^32,48^. Astrocytes were immunolabelled with DAPI and GFAP, and fluorescence images were subdivided into non-overlapping sub-regions (hereafter referred to as “crops”), yielding more than 19,000 image crops spanning a broad spectrum of morphologies, from flattened and spread cells to highly filamentous configurations (**Fig. 1B** and **Table S2**). This morphological diversity reflects the intrinsic heterogeneity of astrocytes in culture^49^ and highlights the analytical challenges posed by dense microscopy datasets with frequent cell overlap^50,51^. To enable direct comparison between imaging-derived and molecular phenotypes, we additionally performed matched whole-transcriptome RNA sequencing across all experimental conditions (**Table S3**).

To systematically characterise cellular phenotypes from microscopy images without relying on segmentation or manual annotation, we trained a ResNet18 model using a modified version of the SimCLR^52^ self-supervised contrastive learning framework. In standard SimCLR, positive pairs are generated by applying independent augmentations to the same image, an approach originally developed for visually diverse natural image datasets^53^. However, biological microscopy images frequently exhibit high inter-sample similarity, increasing the risk that models focus on trivial visual correlations or technical variation rather than biologically meaningful structure^16–18,54,55^. To address this limitation, we introduced spatially neighbouring image crops as additional positive pairs during contrastive training (**Fig. 1C**). Because adjacent crops share local biological context while remaining visually distinct, this strategy increases positive-pair diversity while preserving information related to cellular organisation, neighbourhood structure and multicellular morphology^17,45,46^. The resulting framework, termed SI-SimCLR (Spatially Informed SimCLR), encourages the model to learn context-aware and biologically informative representations while discarding nuisance variation^16–18,45,56,57^.

Building on these learned representations, we developed a framework to infer plausible morphological change dynamics directly from static imaging data, summarised in a “morphological landscape graph” (**Fig. 1D**). Unsupervised clustering was first utilised to identify recurrent morphological substates as discrete regions of the continuous embedding space, which define the nodes of the graph. We then applied a one-versus-all partial optimal transport framework to determine, for each source state, the subset of states that are most geometrically accessible within the embedding space, and the proportions in which they are accessible. This procedure estimates how cells from a given substate can be optimally redistributed across neighbouring substates while minimising transport cost, thereby defining accessibility relationships directly from embedding-space geometry. The identified accessible substates and their associated transport costs, which quantify the magnitude of morphological change required to move between configurations, were used to construct weighted directed edges in the graph. Analysis of the resulting connectivity structure enabled the classification of substates as attractor-like, progenitor-like, transient or highly plastic configurations, providing a topology-aware view of cellular organisation and potential morphological change within the landscape.

Together, these components establish an integrated strategy for reconstructing topology-aware cellular morphological landscapes from dense microscopy data and systematically characterising substate organisation across disease and inflammatory conditions.

### Spatially informed contrastive learning enables robust representation of astrocyte morphology in fluorescence microscopy images

A central requirement for reconstructing cellular phenotypic landscapes is the ability to learn image representations that preserve biologically meaningful morphological variation while remaining robust to technical and experimental heterogeneity. This challenge is particularly pronounced in fluorescence microscopy datasets, where high inter-sample similarity, dense cellular organisation and batch-dependent variation can dominate representation space and obscure subtle biological signals. We therefore first evaluated whether incorporating spatial context during contrastive training improves the ability of self-supervised learning to capture biologically relevant astrocyte morphology.

To benchmark representation quality, we compared SI-SimCLR embeddings against those generated by standard SimCLR^52^, an ImageNet-pretrained ResNet18, two Vision Transformer^58^ models pretrained on CellPainting datasets, and a handcrafted baseline composed of 189 morphology and texture features (**Fig. 2A** and **Table S4**). Visual inspection of Uniform Manifold Approximation and Projection (UMAP) embeddings^59^, coloured by disease state, inflammatory condition and experimental batch, revealed that all methods captured some degree of batch-associated structure, although this effect was less pronounced for the Vision Transformer models (**Fig. 2A**). To quantify the extent to which embeddings encode batch-related variation, we clustered each representation space and computed the Adjusted Rand Index (ARI)^60^ between cluster assignments and batch labels (see *Methods*). While all methods yielded positive ARI scores, SI-SimCLR and, to a lesser extent, the ImageNet-pretrained ResNet18 exhibited the strongest batch-associated structure (**Fig. 2B**).

**Figure 2.**
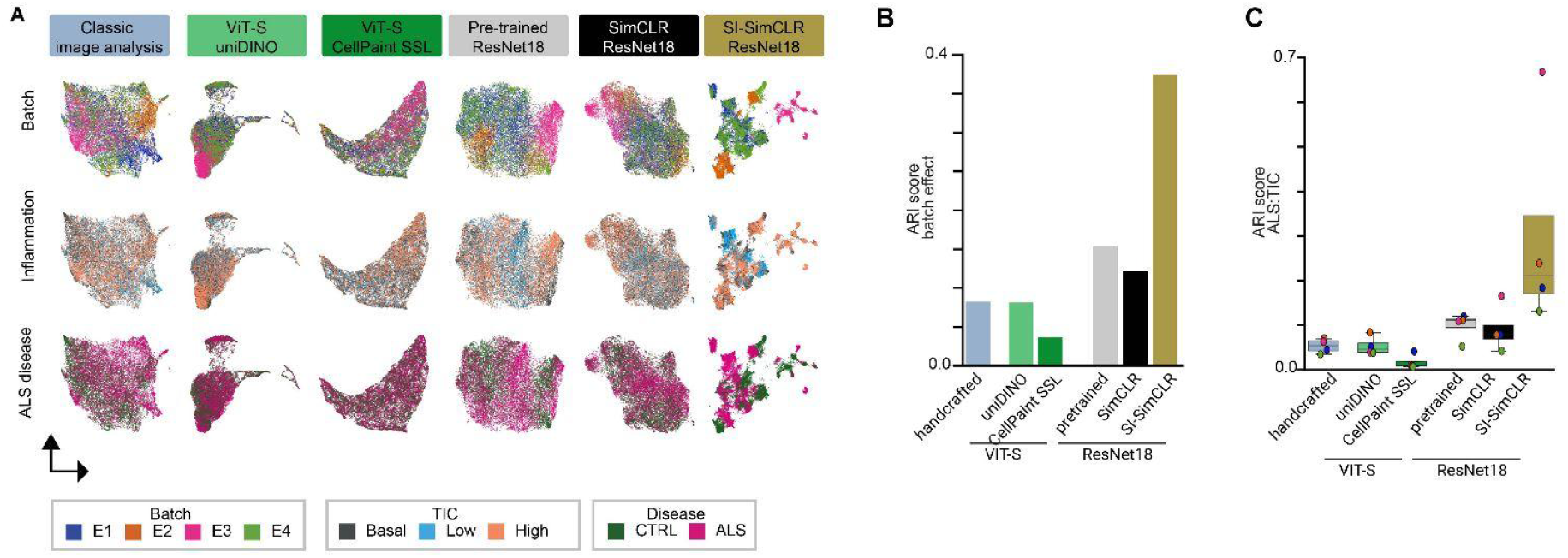
Biologically relevant image representations enabled by spatially informed contrastive learning. **A.** UMAP^99^ visualisation of crop-level embeddings (n = 19,326) obtained using six image representation methods. Embeddings were derived from handcrafted features (*grey*), a ResNet18 pretrained on ImageNet (pink), a ResNet18 trained using the standard SimCLR contrastive pairing strategy (blue), two Vision Transformer models pretrained with DINO on Cell Painting data, uniDINO^14^ trained on single-channel images (*light green*) and CellPaint SSL^76^ trained on five-channel images (*dark green*), and a ResNet18 trained using spatially informed contrastive pairing (SI-SimCLR; *gold*). **B**. Bar plots showing for each model the Adjusted Rand Index (ARI) using batch identity as label. The ARI quantifies the extent to which embeddings cluster according to the 4 batches, with higher values indicating stronger batch-associated structure. For each model, we evaluated a range of UMAP hyperparameters (see *Materials and Methods*) before applying KMeans with K=4, and reported the highest score across the tested UMAP parameters. **C**. Boxplots showing the distribution of ARI scores across individual batches when using biological conditions (ALS/control, TIC treatment) as labels. For each model and experimental batch, we independently evaluated a range of UMAP hyperparameters and applied K-means with K=4 or K=6, depending on the number of conditions in the batch, and reported the highest ARI across the tested UMAP hyperparameters.

To assess whether this batch structure reflects genuine biological variation rather than technical artefact, we first checked that batch labels were not predictive of biological signals with an ARI of 1.8%. Then, we examined the average values of 189 handcrafted features across batches (**Supplementary Fig. 1, Table S4**). Systematic, feature-category-specific differences were apparent: batch E3 exhibited elevated Gabor filter features alongside reduced Haralick texture and image statistics values relative to other batches, while batches E1 and E4 showed the most similar feature profiles overall. Notably, the strongest batch-associated variation was observed in Gabor and Haralick features, which encode local texture structure and spatial intensity dependencies, rather than in global intensity statistics such as mean pixel intensity or Otsu-threshold variance. Although such features can still be influenced by imaging conditions, this pattern is more consistent with differences in cellular morphology and spatial organisation. Purely technical effects, such as staining intensity or illumination variability, would be expected to primarily affect global intensity measures. Together, these observations suggest that the batch variation captured by the embeddings is at least partly driven by underlying biological heterogeneity in astrocyte morphology across independent culture preparations.

Importantly, SI-SimCLR embeddings were the only ones in which biological signals were visually distinguishable across all conditions in UMAP space (**Fig. 2A**). To quantify biological signals independently of batch effects, we performed unsupervised clustering on each batch separately and measured the correspondence between cluster assignments and biological conditions (ALS versus control, basal versus TIC-low versus TIC-high) using the ARI. While all methods yielded modest positive scores, SI-SimCLR consistently achieved the highest within-batch ARI across batches (ranging from 13% to 67%), indicating superior capture of biologically relevant variation (**Fig. 2C**). To assess whether these biological signals identified within individual batches generalise across experiments, we applied a leave-one-batch-out strategy. MLP classifiers were trained on three batches and evaluated on the held-out batch to distinguish ALS from control and TIC-treated from untreated astrocytes. Across all classifiers and held-out batches, SI-SimCLR embeddings consistently achieved the highest AUC, demonstrating that the captured biological signals are reproducible and transferable rather than batch-specific (**Supplementary Figs. 2A,B**). Batches E1 and E4 showed the most consistent cross-batch classification performance, suggesting that their embeddings are less dominated by batch-specific morphological variation and therefore provide the most reliable foundation for cross-batch biological inference.

Together, these results show that incorporating local spatial context during contrastive learning enables SI-SimCLR to recover biologically meaningful morphological organisation that generalises across experimental batches, providing a robust foundation for reconstruction of cellular phenotypic landscapes.

### Topology-aware reconstruction of astrocyte morphological landscapes reveals interconnected and plastic cellular substates

We next constructed a morphological atlas of cultured astrocytes using the biologically informative image representations learned by SI-SimCLR. To maximise sensitivity to condition-associated morphological variation while preserving the global organisation of the embedding space, unsupervised clustering was performed within batch E1, which displayed the strongest and most reproducible biological signal (**Fig. 2C** and **Supplementary Figs. 2A,B**). This analysis identified 12 dominant astrocyte morphological substates containing variable numbers of image crops (**Figs. 3A–C;** see *Methods*). Collectively, these substates captured substantial phenotypic diversity, spanning morphologies from rounded and spread configurations to highly filamentous structures (**Fig. 3D**), consistent with the pronounced morphological plasticity previously described in astrocytes^61,62^.

**Figure 3.**
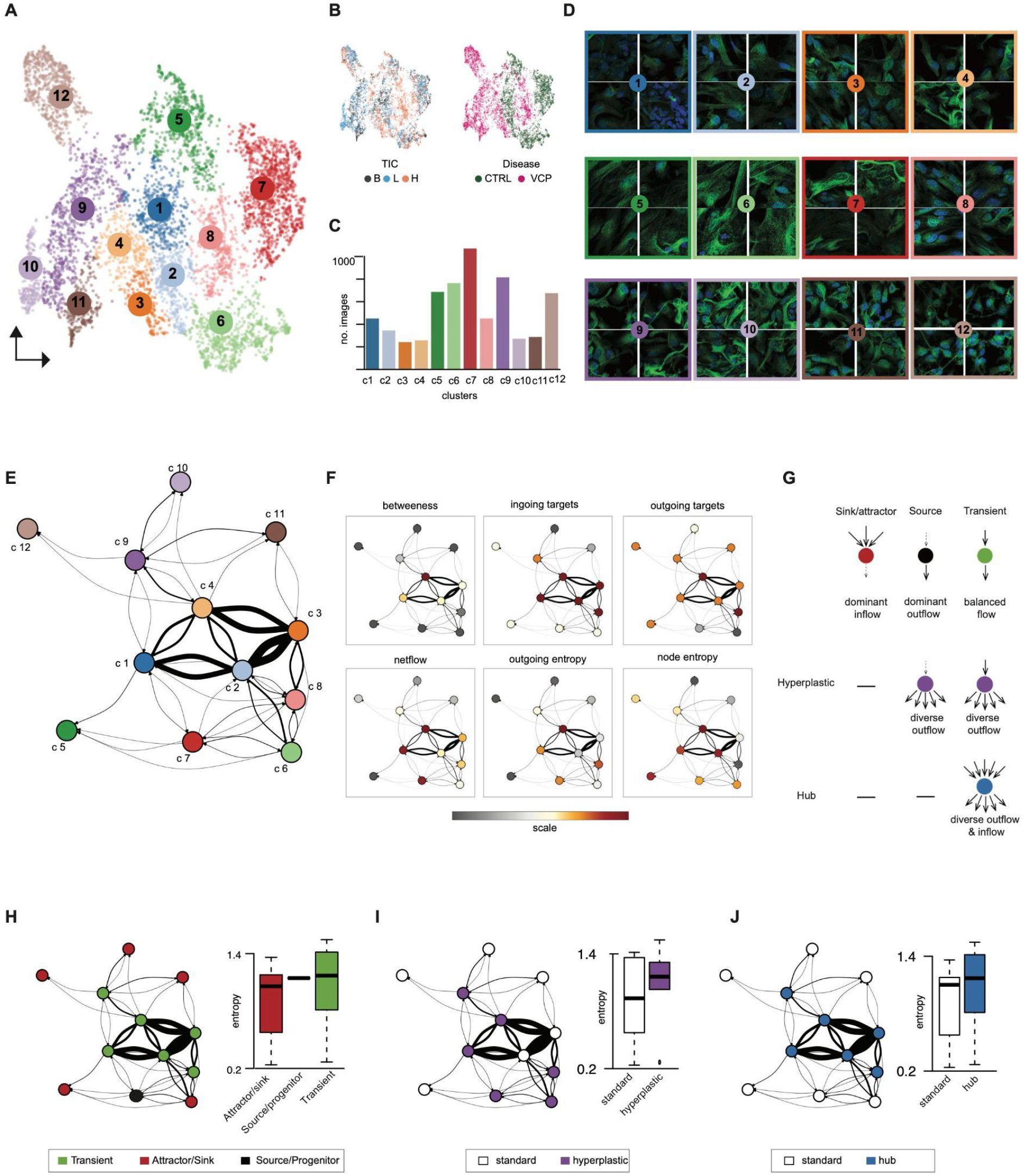
A morphological atlas of astrocyte substates reveals structured phenotypic heterogeneity and state transition topology across ALS and neuroinflammatory conditions. **A.** UMAP^90,9999^ visualisation of crop-level SI-SimCLR embeddings from experimental batch E1 (n = 6,321 crops), coloured by the 12 morphologically distinct substates identified by Shared Nearest Neighbour (SNN)^90^ unsupervised clustering. **B**. Same UMAP as (A), coloured by TIC inflammatory condition (*left*; B: basal, L: low, H: high) and disease status (*right*; CTRL: control, VCP: VCP-mutant ALS). **C**. Bar chart showing the number of image crops per cluster, reflecting the variable size of morphological substates across the atlas. **D**. Representative image crops for each of the 12 clusters, with cluster identity indicated by coloured borders matching (A). The four crops closest to the cluster centroid are shown, illustrating the internal coherence of each substate and the morphological diversity across the atlas, spanning a spectrum from rounded and spread configurations to elaborately filamentous arrangements. **E**. Morphological transition graph in which nodes represent clusters and directed edges reflect the relative probability of phenotypic interconversion between substates, estimated via optimal transport (OT) over SI-SimCLR embedding distributions. Edge thickness and colour intensity are proportional to transition probability; edges below the Q3 transition probability threshold are pruned for visual clarity. Nodes are coloured by cluster identity. **F**. Transition graphs with nodes coloured by six graph-theoretic metrics: betweenness centrality, number of ingoing targets, number of outgoing targets, net flow, Shannon entropy of outgoing transition flux (outgoing entropy), and biological condition Shannon entropy (node entropy). All metrics are displayed on a continuous grey-to-orange-to-red scale, with darker and warmer colours indicating higher values. **G**. Schematic illustrating the classification criteria for functional state types, Sink/Attractor, Source, Transient, Hyperplastic, and Hub, based on their characteristic inflow, outflow, and entropy profiles. Sink/Attractor states are defined by dominant inflow; Source states by dominant outflow; Transient states by balanced inflow and outflow; Hyperplastic states by diverse outflow from Source or Transient configurations; and Hub states by both diverse inflow and outflow. **H**. Transition graph with nodes coloured by functional classification, Transient (green), Attractor/Sink (red), and Source/Progenitor (black), alongside a boxplot comparing biological condition Shannon entropy across the three classifications, demonstrating that Transient states exhibit higher condition entropy than Attractor/Sink or Source/Progenitor states. **I–J**. As in (H), with nodes reclassified as standard versus hyperplastic (I; white vs. purple) and standard versus hub (J; white vs. blue). In both cases, hyperplastic and hub substates exhibit significantly higher biological condition entropy than standard substates, consistent with their characterisation as morphological states accessible across diverse biological conditions.

We next sought to determine how these substates are organised relative to one another within the broader morphological landscape. Because the data consist of static imaging snapshots rather than time-resolved measurements, direct state transitions cannot be observed experimentally. We therefore developed a graph-based framework to infer plausible relationships between substates directly from the geometry of the SI-SimCLR embedding space (**Fig. 3E**; *Methods*). In this framework, each node represents a morphological substate identified by clustering, whereas directed edges represent inferred transition accessibility between substates estimated using partial optimal transport (OT). Edge weights quantify the relative ease with which one morphological configuration can be transformed into another within the learned representation space.

Unlike pairwise distance-based approaches, the OT framework evaluates how each source substate relates simultaneously to all other substates, thereby identifying sparse and geometry-constrained accessibility patterns across the landscape. The resulting graph therefore represents the global connectivity structure of the morphological landscape rather than direct experimentally observed trajectories. Conceptually, this organisation approximates a Waddington-like landscape^20^ reconstructed from static imaging data, in which stable configurations behave as attractor-like states, whereas intermediary regions connect multiple phenotypic configurations.

To characterise the organisation of this landscape, we quantified several graph-theoretic properties of each substate, including weighted incoming connectivity (*in-strength*), weighted outgoing connectivity (*out-strength*), outgoing transition entropy and betweenness centrality (**Fig. 3F**). Betweenness centrality measures how frequently a substate lies along highly connected paths linking distinct regions of the graph and therefore identifies intermediary configurations connecting otherwise separated landscape regions. Outgoing transition entropy quantifies how broadly a substate distributes its outgoing accessibility across downstream states. High entropy indicates that a state is connected to many alternative downstream configurations rather than preferentially linked to a single dominant destination.

Using these properties, we classified substates into three principal topological archetypes (**Fig. 3G**): *source/progenitor* states characterised by dominant outgoing accessibility; *attractor/sink* states characterised by dominant incoming accessibility; and *transient* states characterised by elevated betweenness centrality and intermediate positioning within the graph. This analysis identified a single source/progenitor state (c7), whereas the remaining substates distributed across attractor/sink and transient classes (**Fig. 3H**). Consistent with these inferred roles, transient substates occupied central graph positions connecting multiple regions of the landscape, whereas attractor/sink states preferentially localised to peripheral regions.

We further defined high-plasticity states as substates combining high outgoing transition entropy with broad downstream connectivity. Importantly, high-plasticity in this context does not refer to experimentally observed temporal plasticity, but rather to the inferred accessibility of multiple alternative morphological configurations within the reconstructed landscape. Most high-plasticity substates overlapped with transient substates, indicating that central regions of the landscape connect broadly across multiple phenotypic configurations rather than through narrow deterministic paths. The sole source/progenitor state c7 also displayed high outgoing entropy despite its peripheral graph position, whereas transient states c2 and c3 were not classified as high-plasticity states because their outgoing accessibility remained concentrated toward a restricted subset of substates (**Fig. 3I**). Hub states, defined by extensive connectivity to both upstream and downstream regions of the graph, overlapped strongly with transient substates (**Fig. 3J**), indicating that intermediary regions of the astrocyte morphological landscape are highly interconnected rather than organised as isolated linear bridges.

To evaluate the biological relevance of the inferred landscape topology, we next asked whether highly connected substates preferentially integrated cells originating from diverse biological conditions. We reasoned that if graph connectivity reflects biologically meaningful accessibility relationships, highly connected substates should be shared across multiple disease and inflammatory contexts rather than restricted to a single condition. We therefore quantified the distribution of image crops originating from each biological condition within every substate (ALS versus control under basal, TIC-low and TIC-high exposure; **Supplementary Fig. 3A**) and computed Shannon entropy as a measure of condition heterogeneity within each node. In this context, high condition entropy indicates that a substate contains cells originating from diverse biological conditions, whereas low condition entropy reflects condition-restricted occupancy.

High-plasticity, transient and hub substates consistently displayed elevated condition entropy relative to source/progenitor and attractor/sink states (**Figs. 3H–J**). Thus, substates inferred to occupy highly connected regions of the graph were also preferentially populated by cells from diverse biological contexts. The convergence between graph topology and condition heterogeneity supports the biological relevance of the reconstructed landscape organisation and suggests that highly connected substates correspond to morphologically accessible intermediary configurations shared across multiple cellular contexts.

Together, these analyses establish a topology-aware morphological atlas composed of source-like, attractor-like and highly interconnected intermediary substates. More broadly, this framework provides a principled and condition-agnostic strategy for reconstructing continuous cellular state organisation from dense microscopy data and for investigating how disease and environmental perturbations reshape phenotypic landscape structure.

### ALS-associated *VCP* mutation defines a distinct astrocyte morphological landscape remodelled by neuroinflammation

Having established the topology of the astrocyte morphological landscape, we next investigated how disease genotype and inflammatory stimulation shape substate occupancy across the graph. Although most substates were co-occupied by multiple biological conditions, we observed a pronounced genotype-dependent organisation of the landscape. The majority of substates were significantly enriched for either control or VCP-mutant astrocytes, largely independently of TIC exposure, indicating that disease genotype exerts a dominant influence on astrocyte morphological identity that supersedes inflammatory context (**Supplementary Fig. 3B**). Under basal conditions, control and *VCP*-mutant astrocytes occupied largely distinct regions of the landscape. Control astrocytes were preferentially enriched in substates c1, c2, c5, c6, and c7, characterised by spread and filament-rich morphologies, whereas *VCP*-mutant astrocytes preferentially occupied substates c4, c9, c11, and c12, associated with more elaborate filamentous configurations (**Figs. 4A,B**).

**Figure 4.**
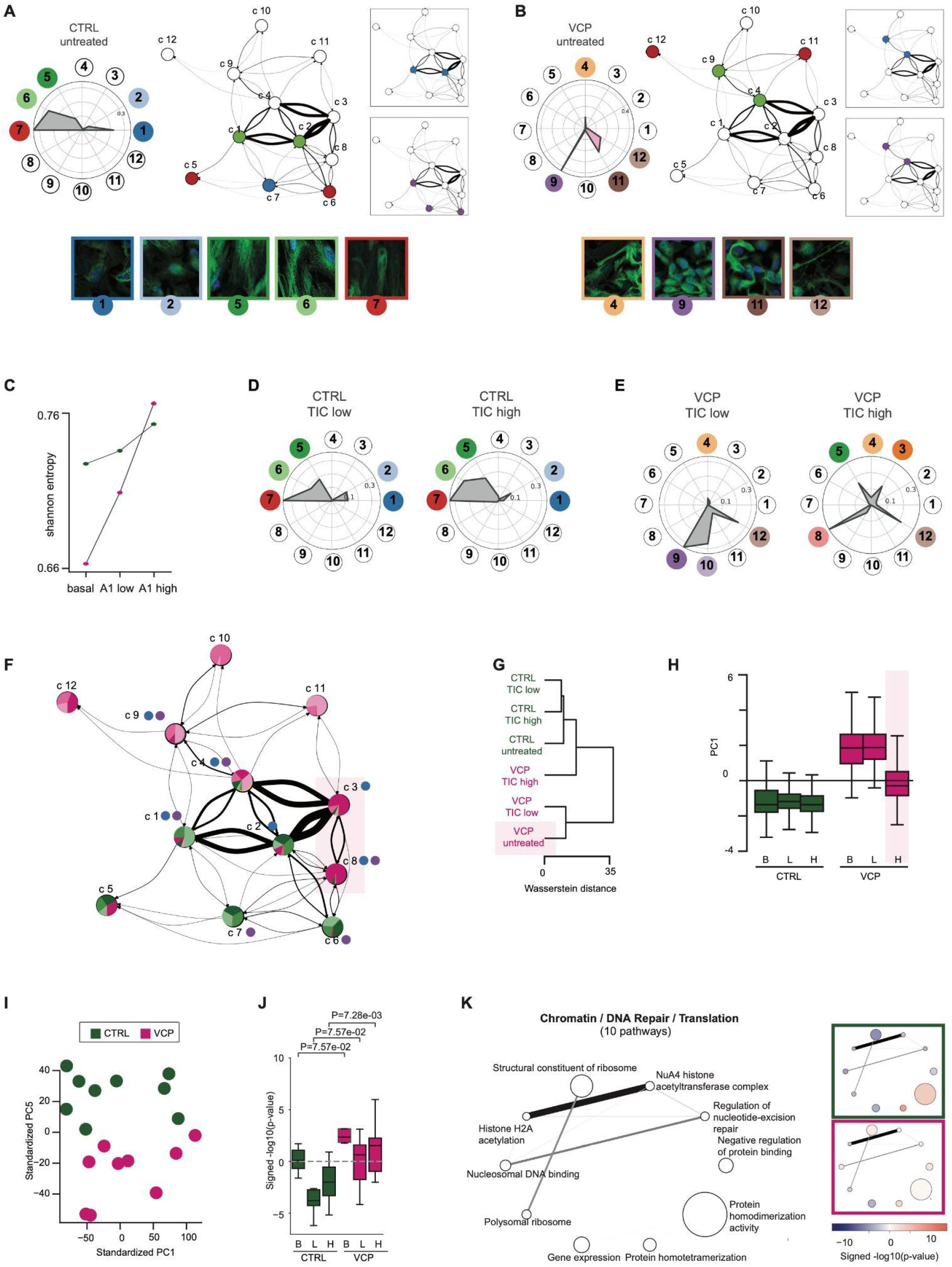
Morphological and transcriptional profiling capture complementary but non-identical aspects of astrocyte state organisation. **A.** Radar plot (*left*) showing the proportional representation of each morphological substate for control astrocytes under basal conditions, with coloured circles indicating cluster identity. The corresponding subgraph (*right*) displays the active transition structure among control-enriched substates, with edge thickness proportional to transition probability. Representative image crops for the five most enriched control substates are shown below, with cluster identity indicated by coloured borders. **B**. As in (A) for VCP-mutant astrocytes under basal conditions. The radar plot and subgraph highlight the four substates most enriched in VCP-mutant astrocytes, which occupy a distinct and more peripheral region of the transition graph relative to control-enriched substates. **C**. Shannon entropy of cluster occupancy as a function of inflammatory condition (basal, TIC-low, TIC-high) for control (*green*) and VCP-mutant (*magenta*) astrocytes. Each point represents the entropy of the distribution of images across the 12 morphological substates for a given genotype and condition. Higher entropy indicates greater dispersal across the morphological landscape. **D**. Radar plots showing proportional substate occupancy for control astrocytes under TIC-low (*left*) and TIC-high (*right*) inflammatory conditions. Coloured circles indicate cluster identity as in (A). **E**. As in (D) for VCP-mutant astrocytes under TIC-low (left) and TIC-high (right) conditions, showing a pronounced redistribution of cells across substates under high inflammatory stimulation relative to basal conditions. **F**. Morphological transition graph with nodes coloured by genotype composition (green: control-enriched; magenta: VCP-enriched) and treatment (light shades: untreated; medium shades: TIC-low; dark shades: TIC-high). Edge thickness is proportional to transition probability. The light pink box highlights the region of the graph where VCP-enriched substates selectively expanded under TIC-high conditions (c3 and c8) connect to control-enriched regions of the graph. **G**. Dendrogram showing hierarchical clustering of the six biological conditions based on Wasserstein distances between SI-SimCLR embedding distributions. Ward’s linkage was used. Conditions are coloured by genotype (green: control; magenta: VCP-mutant), demonstrating that genotype is the primary driver of morphological separation: all control conditions group together, and all VCP conditions form a distinct clade. VCP TIC-high is positioned closest to the boundary with control conditions. **H**. Scatter plot of the first two principal components of SI-SimCLR crop embeddings, coloured by genotype (green: control; magenta: VCP-mutant). PC1 (18% variance explained) separates the two genotypes, with VCP TIC-high astrocytes showing reduced separation along this axis relative to VCP basal and TIC-low conditions. **I**. Scatter plot of standardised PC1 (17% variance explained) and PC5 (8.2% variance explained) projections from gene expression data. Each point represents one sample, coloured by genotype. PC5 captures the transcriptional contribution of the VCP mutation, which is subtler than the treatment-driven variation captured by PC1. **J**. Box plots showing inferred activity scores for 10 biological pathways related to chromatin organisation, DNA repair, and translation, selected based on their contribution to the VCP-associated principal component of pathway activity (see Materials and Methods). Samples are grouped by genotype and inflammatory condition (B: basal; L: TIC-low; H: TIC-high). Statistical comparisons between control and VCP-mutant astrocytes are indicated with *P*-values. Boxes indicate median and interquartile range; whiskers extend to 1.5× the interquartile range. **K**. Network representation of the 10 pathways selected in (K). Nodes correspond to Gene Ontology pathways and are sized proportionally to the number of annotated genes. Edges indicate gene overlap between pathways quantified by the Jaccard similarity index (shown when index > 0.01); edge width scales with similarity. Node colours represent pathway enrichment scores on a diverging scale (red: enriched; blue: depleted) shown separately for control (green border) and VCP-mutant (magenta border) astrocytes under TIC-low conditions.

To quantify phenotypic heterogeneity, we computed the Shannon entropy of substate occupancy distributions for each condition. High entropy reflects broad occupancy across multiple substates, whereas low entropy indicates restriction to a limited subset of morphologies. Under basal conditions, *VCP*-mutant astrocytes exhibited lower entropy than controls, indicating that ALS-associated astrocytes occupy a more constrained morphological state space at baseline (**Fig. 4C**). Inflammatory stimulation increased phenotypic heterogeneity in both genotypes, but the effect was markedly more pronounced in *VCP*-mutant astrocytes, with TIC-high conditions leading to the strongest redistribution across the landscape. The latter is characterised by reduced occupancy of the dominant *VCP*-associated cluster c9 together with increased representation of intermediary and control-associated substates, whereas control astrocytes exhibited comparatively modest redistribution following TIC exposure (**Figs. 4D,E**).

Visualisation of the morphological landscape graph confirmed that control- and *VCP*-enriched substates occupy largely segregated regions connected through a limited number of intermediary substates (**Fig. 4F**). Notably, clusters c3 and c8, which consisted mainly of *VCP*-mutant astrocytes under TIC-high conditions, corresponded to highly connected hub-like substates bridging control- and *VCP*-associated regions of the graph, suggesting that inflammatory remodelling of ALS astrocytes preferentially occurs through broadly accessible intermediary configurations rather than direct transitions between genotype-restricted states. This organisation was independently supported by Wasserstein-distance hierarchical clustering and principal component analysis of embedding distributions. Biological conditions clustered primarily by genotype, with all control conditions grouping separately from *VCP*-mutant conditions (**Figs. 4G,H**). Nevertheless, *VCP* TIC-high astrocytes consistently shifted toward the control-associated region of embedding space, consistent with the entropy and graph-topology analyses.

To determine whether the ALS-dependent organisation observed in the morphological landscape was similarly reflected at the transcriptional level, we analysed bulk RNA-seq data across the same conditions. In contrast to the morphology-based analyses, transcriptome-wide organisation was driven primarily by inflammatory stimulation rather than genotype. Both hierarchical clustering based on Spearman correlation and principal component analysis revealed strong treatment-associated separation, with PC1 (17% variance explained) capturing the overall inflammatory response independently of TIC dose or duration, and PC7 (6.4%) distinguishing TIC-low from TIC-high conditions (**Supplementary Figs. 4A–D**). By comparison, the contribution of the VCP mutation to global transcriptional variation was more modest and captured primarily by PC5 (8.2% variance explained; **Fig. 4I**). Pathway activity analysis revealed that inflammatory stimulation perturbed largely overlapping biological processes in both genotypes, including reduced cell adhesion pathway activity, consistent with cytoskeletal remodelling and altered cell–cell interactions, together with increased activation of stress and immune response programmes (**Supplementary Fig. 4E**). Nevertheless, VCP-specific transcriptional signatures remained detectable at the pathway level, including increased activity of DNA repair pathways and dysregulation of cytoskeletal remodelling programmes selectively enriched in VCP-mutant astrocytes (**Figs. 4J,K** and **Supplementary Fig. 4F**).

Together, these findings show that morphological and transcriptional profiling capture complementary but non-identical aspects of astrocyte state organisation. Whereas inflammatory signalling dominated global transcriptional variation, astrocyte morphology remained primarily structured by ALS-associated VCP genotype. Although VCP-dependent signatures were detectable in both modalities, disease-associated morphological states encoded an additional layer of cellular organisation not fully resolved by transcriptomic programmes alone. Under basal conditions, VCP-mutant astrocytes occupied a constrained and genotype-specific region of the morphological landscape that partially redistributed toward control-associated configurations following strong neuroinflammatory stimulation.

### Distinct *VCP*-mutant astrocyte substates exhibit differential inflammatory reactivity

We previously showed that *VCP*-mutant ALS astrocytes adopt diverse cell-autonomous reactive states at the transcriptomic level^33^. We therefore asked whether the morphological substates identified here similarly reflect distinct reactive configurations and whether individual *VCP*-mutant substates differ in their relationship to inflammatory astrocyte states. To address this, we used partial OT to quantify the similarity between untreated *VCP*-mutant astrocyte substates and control astrocytes under basal, TIC-low, or TIC-high inflammatory conditions. This analysis revealed marked substate-specific differences in inflammatory affinity. The dominant *VCP*-mutant substate c9, a transient-like configuration accounting for the majority of untreated *VCP*-mutant astrocytes, exhibited the strongest similarity to untreated control astrocytes in line with its spread morphology (**Figs. 5A–D**). In contrast, substates c11 and c12, both attractor/sink states characterised by denser, more compact morphologies, displayed progressively stronger affinities for TIC-low and TIC-high inflammatory conditions, respectively (**Figs. 5C–E**). Cluster c4, a high-plasticity hub-like substate, exhibited intermediate affinities across conditions, consistent with its intermediary position within the morphological landscape.

**Figure 5.**
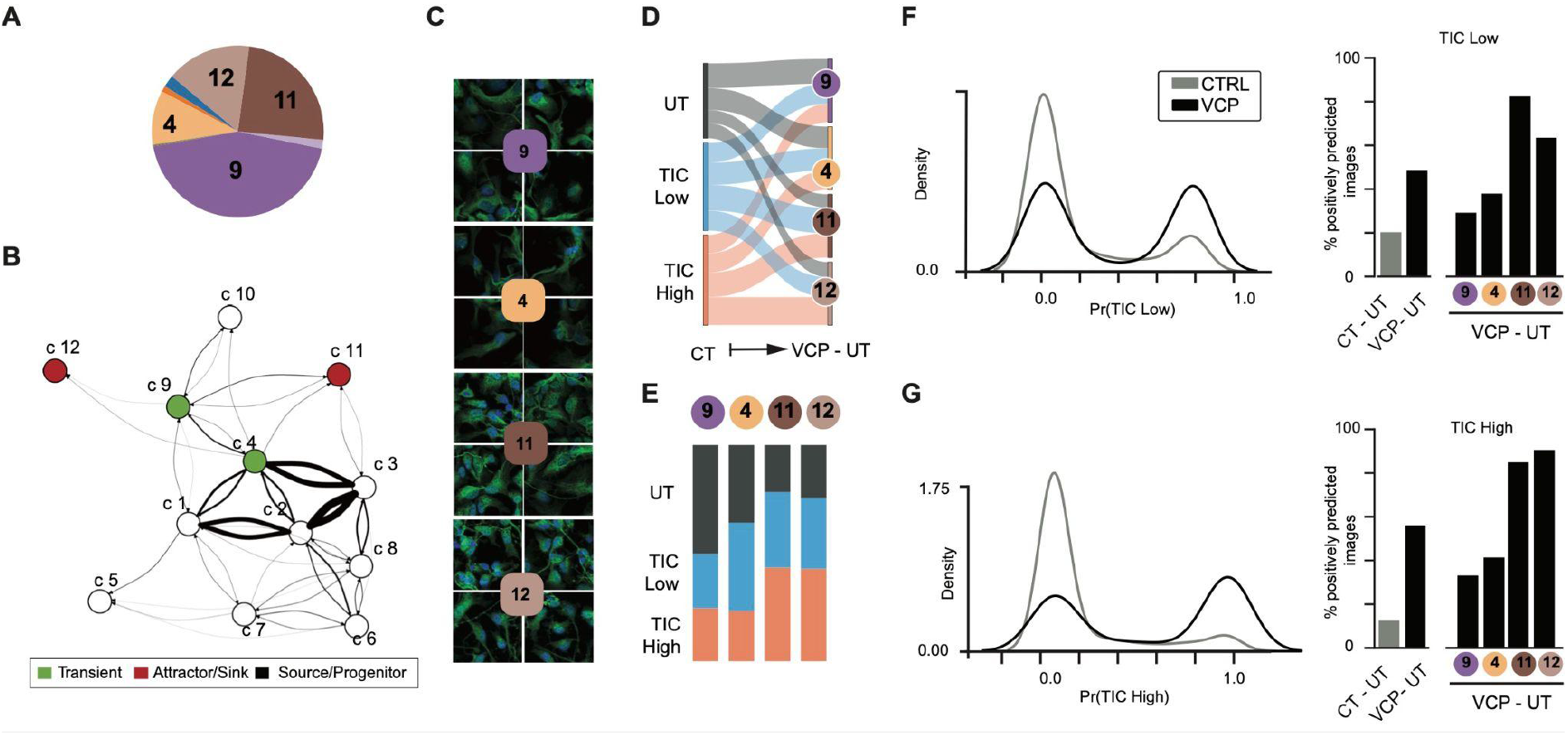
VCP-mutant astrocytes harbour morphologically heterogeneous substates with distinct relationships to inflammation-induced reactive configurations. **A.** Pie chart showing the proportional size of the four VCP-enriched morphological substates under basal conditions (c9, c11, c12, c4), with c9 constituting the dominant substate. **B**. Subgraph showing the transition structure among the four VCP basal substates within the full morphological transition graph. Node colour indicates functional classification: hub/transient (green, c4) and source/progenitor (blue, c9, c11, c12). Edge thickness is proportional to transition probability. **C**. Representative brightfield microscopy images of astrocytes belonging to each of the four VCP basal substates (c9, c4, c11, c12), illustrating their distinct morphological identities. **D**. Alluvial (Sankey) diagram representing optimal transport plans from control reactive conditions (untreated control [CT], TIC-low, TIC-high) to the four VCP basal substates. Transport was computed using partial optimal transport and normalised per VCP substate. Flow width encodes the magnitude of the optimal coupling, indicating the relative morphological affinity between each control condition and each VCP substate. **E**. Stacked bar charts showing the proportional contribution of each control condition (dark grey: untreated; blue: TIC-low; orange: TIC-high) to each of the four VCP basal substates (c9, c4, c11, c12), derived from the optimal transport plans in (D). Cluster c9 receives predominantly from untreated control, whereas c11 and c12 show proportionally greater affinities toward TIC-low and TIC-high reactive conditions, respectively, revealing substate-specific relationships to inflammatory reactivity within the VCP basal population. **F**. Density distributions of predicted probabilities from an MLP classifier trained to distinguish untreated control from TIC-low reactive astrocytes (*left*), applied to untreated control (grey) and VCP-mutant (black) astrocytes. VCP-mutant astrocytes display a pronounced secondary peak near Pr = 1.0, absent in controls, indicating that a substantial subpopulation of untreated VCP cells is assigned a high probability of TIC-low reactive identity. The bar chart (*right*) shows the percentage of images from control (CT-UT, VCP-UT) and each VCP basal substate (c9, c4, c11, c12) classified as positive (probability threshold = 0.5) for TIC-low reactivity, revealing heterogeneous classifier responses across substates. **G**. As in (F) for the TIC-high classifier. Approximately 55% of untreated VCP-mutant images are classified as TIC-high reactive, compared with near zero in untreated controls. Bar chart (*right*) shows substate-level variation in TIC-high classification rates, with c11 and c12 contributing disproportionately to the TIC-high-like signal within the VCP basal population.

To independently validate these observations, we trained multilayer perceptron (MLP) classifiers on SI-SimCLR embeddings to distinguish TIC-low and TIC-high reactive astrocytes from untreated controls, and applied these classifiers to untreated *VCP*-mutant astrocytes. Consistent with the optimal transport analysis, untreated *VCP*-mutant astrocytes showed substantially increased predicted inflammatory reactivity relative to untreated controls, with approximately 45% and 55% of images assigned high TIC-low and TIC-high probabilities, respectively (**Figs. 5F,G**, *left*). Importantly, this inflammatory resemblance was highly substate-dependent. Cluster c9 showed the weakest inflammatory association, whereas c11 and c12 displayed the strongest overlap with TIC-low and TIC-high reactive states, respectively (**Figs. 5F,G**, *right*). Conversely, TIC-treated control astrocytes were only rarely classified as ALS by an ALS-versus-control classifier (**Supplementary Fig. 5**), indicating that inflammatory stimulation alone does not fully recapitulate the ALS morphological phenotype.

Taken together, these findings demonstrate that untreated VCP-mutant astrocytes harbour heterogeneous cell-autonomous morphological states that differentially recapitulate inflammation-associated reactive configurations. Importantly, these reactive affinities are structured at the substate level rather than uniformly distributed across the population, underscoring the importance of substate-resolved analysis for disentangling complex disease phenotypes. More broadly, these results illustrate how the atlas framework enables decomposition of ALS-associated astrocyte heterogeneity into interpretable morphological programmes linked to distinct inflammatory reactive states.

## DISCUSSION

We present SI-SimCLR, a spatially informed self-supervised learning framework that extracts biologically informative representations from high-content fluorescence microscopy images without requiring segmentation or predefined labels. Applied to a large multimodal dataset of human iPSC-derived astrocytes spanning ALS genotype and inflammatory conditions, SI-SimCLR enabled reconstruction of a topology-aware morphological atlas capturing continuous astrocyte state organisation across biological contexts. Unlike handcrafted descriptors, which failed to robustly resolve disease- and inflammation-associated variation, SI-SimCLR recovered reproducible morphological structure across experimental batches, indicating that relevant pathological information is encoded in complex spatial and multicellular features not readily captured by predefined morphological measurements.

A central contribution of this work is the combination of representation learning with graph-based landscape reconstruction. Most imaging analyses treat cellular states as isolated categories or compare populations using global summary statistics. In contrast, our framework models morphology as a continuous and structured landscape in which substates are interconnected through inferred accessibility relationships derived from the geometry of the embedding space. This graph representation provides information not only about which substates exist, but also about how they are organised relative to one another, including which configurations behave as stable attractor-like states, which occupy intermediary positions, and which remain broadly connected across multiple phenotypic regions. The convergence between graph connectivity and biological-condition heterogeneity further supports the biological relevance of the inferred topology, suggesting that highly connected substates correspond to morphologically accessible and plastic cellular configurations.

Within this landscape, ALS-associated VCP mutation strongly constrained astrocyte morphological organisation under basal conditions, with mutant astrocytes preferentially occupying a restricted genotype-associated region of the landscape. Inflammatory stimulation increased morphological heterogeneity in both genotypes, but produced substantially greater redistribution in VCP-mutant astrocytes, shifting cells toward intermediary and control-associated substates. Substate-level analyses further revealed that untreated VCP-mutant cultures are themselves heterogeneous and contain multiple partially reactive morphological configurations. Together, these findings suggest that ALS-associated astrocyte states do not represent fixed phenotypic endpoints, but instead occupy structured regions within a broader and dynamically accessible morphological landscape.

A second major finding is that morphological and transcriptomic profiling capture complementary but non-identical aspects of astrocyte state organisation. Whereas inflammatory signalling dominated global transcriptional variation, astrocyte morphology remained primarily structured by ALS-associated VCP genotype. Although VCP-dependent signatures were detectable in both modalities, their relative contributions differed substantially, indicating that imaging-derived morphology captures an additional layer of cellular organisation not fully resolved by dominant transcriptional programmes alone. In this context, imaging-based phenotyping may provide a particularly powerful readout of disease-associated cellular organisation in systems where pathological states are spatially heterogeneous, transient or incompletely reflected by population-averaged molecular measurements.

An important technical consideration concerns the biological interpretation of batch structure captured by SI-SimCLR embeddings. Unlike histopathology datasets, where batch effects often predominantly reflect technical artefacts, differences between iPSC-derived astrocyte culture batches may themselves encode biologically meaningful variation related to cell density, passage history or local culture state. The sensitivity of SI-SimCLR to such variation may therefore reflect its ability to capture subtle biological organisation rather than merely technical confounding. Nevertheless, disentangling biologically meaningful variability from unwanted batch-specific effects remains an important challenge for future self-supervised imaging frameworks.

Several limitations should also be acknowledged. Although the present study focused on ALS-associated VCP mutation, extending these analyses across broader genetic backgrounds, primary human astrocytes and in vivo systems will be important to establish the generalisability of the landscape organisation described here. Similarly, incorporating additional astrocyte markers beyond GFAP may further refine phenotypic resolution and improve characterisation of reactive and homeostatic substate diversity. More broadly, this work demonstrates how combining self-supervised representation learning with topology-aware graph analysis enables reconstruction of continuous cellular state landscapes directly from microscopy data. Because this framework does not require segmentation, predefined labels or experimentally observed trajectories, it is broadly applicable to dense imaging systems in which cellular organisation itself is a key component of phenotype. Beyond disease modelling, this approach may provide a powerful framework for phenotypic screening by enabling quantitative analysis not only of state occupancy, but also of how genetic or pharmacological perturbations reshape landscape connectivity, plasticity and transition accessibility across cellular populations.

## MATERIAL AND METHODS

### hiPSC astrocyte culture

We used our previously published differentiation paradigm to generate highly enriched (>90%) populations of hiPSC-derived astrocytes. All cell cultures were maintained at 37°C and 5% carbon dioxide. hiPSCs were cultured on Geltrex (Thermo Fisher Scientific) coated plates in Essential 8 medium (Thermo Fisher Scientific) and routinely passaged with EDTA. To generate hiPSC-derived astrocytes, we employed a modified version of the protocol established by Hall et al. 2017. Following a 7-day neural conversion in chemically defined medium supplemented with 1 μM dorsomorphin (Millipore), 2 μM SB431542 (Tocris Bioscience), and 3.3 μM CHIR99021 (Miltenyi Biotec), precursors were patterned using 0.5 μM retinoic acid and 1 μM purmorphamine for 7 days, followed by a 4-day treatment with 0.1 μM purmorphamine. After a long-term propagation phase (>60 days) with FGF-2 (10 ng/ml), cells underwent terminal differentiation in BMP4 and LIF for 23 days, followed by a 7-day differentiation period in N2B27 alone. The latter was implemented as a modification to a previously published protocol^8^ to ensure the astrocytes reached a more basal, unstimulated state prior to experimental use.

### Pro-inflammatory factors treatment

Astrocytes were treated for low TIC condition with different concentrations of human TNF-ɑ (30 ng/ml; R&D 210-TA/CF), human IL-1ɑ (3 ng/ml; Sigma-Aldrich), and human C1q (400 ng/ml; MyBiosource) for 24 hrs. For TIC high condition, astrocytes were treated with human TNF-ɑ (100 ng/ml), human IL-1ɑ (100ng/ml), human C1q (1μg/ml) for 6 days with media change every 2 days^63,64^. This enables the study of morphological changes associated with cytokine-induced reactive transformation^63,65^

### Immunofluorescence & Imaging

Astrocytes were seeded in 96-well plates at densities of 20,000 cells/well, across 4 different experiments. Prior to immunocytochemistry, cultures were washed with PBS and fixed in 4% paraformaldehyde (Sigma-Aldrich) for 15 minutes. Following three PBS washes, samples were stored at 4°C. For immunolabelling, cells were blocked and permeabilised for one hour using 5% BSA and 0.3% Triton X-100. GFAP (abcam, 1:1000) was applied overnight at 4°C in the same buffer. After three 5-minute PBS washes, cells were incubated with 488-Alexa Fluor secondary antibody (Thermo Fisher, 1:1000) for one hour at room temperature. Nuclei were counterstained with DAPI (100 ng/ml) for 10 minutes, followed by a final series of three 5-minute PBS washes. Images were acquired using the Opera Phenix High-Content Screening System (Perkin Elmer). Images were acquired with a 40× water objective as confocal z-stacks with z-steps of 1 μm.

### RNA extraction & pre-processing for RNA sequencing

Cells were washed with 1x cold PBS, then RNA was extracted using the Maxwell RSC simplyRNA Cells kit (Promega) and the Maxwell RSC 48 machine. Nanodrop was used to measure RNA quantity and purity. The RNA sequencing libraries used in this study were created from polyadenylated RNA only (polyA+), sequenced at high depth (>30 million reads). PolyA+ library generation was carried out by The Francis Crick Institute. NEBNext Ultra II Directional PolyA mRNA kit was used by the Advanced Sequencing Facility (ASF). RNA-seq libraries are from paired-end stranded sequences and were sequenced using Illumina NovaSeq 6000.

### Image pre-processing and filtering

Raw images are 16-bit images. Raw z-stack images (1080*1080 pixels) from the same field of view were first merged using Maximum Intensity Projection (MIP)^66^, where the pixel with maximum intensity across all z-stacks is selected at each location in the image. For each well, 10 to 12 non-overlapping fields of view (FOVs; 1080×1080 pixels, 0.316 μm/px) were acquired at random locations, yielding over 1,000 FOVs (**Supplementary Table S1**). For all images, DAPI and GFAP fluorescent markers were assigned to the blue and green channels, respectively, while the red channel is set to be pitch-black. After converting MIP images to 8-bit, the channels were merged to form an RGB image. Images were then enhanced using Python Image Library Pillow ImageOps autocontrast function, to normalise image contrast. This function calculates a histogram of the input image, removes 0.1% of the lightest and darkest pixels from the histogram, and remaps the image so that the darkest pixel becomes black (0) and the lightest becomes white (255). The enhanced images were then divided into 16 smaller images of size 270*270 pixels. After cropping, we obtain 23,162 crops from 1,448 fields of view (**Supplementary Table S2**).

To remove low-content images, we calculated the Shannon entropy for the DAPI and GFAP channels of each crop (**Supplementary Fig. 6A**). We then clustered the entropy pairs (DAPI entropy, GFAP entropy) using the HDBSCAN algorithm^67^. We set the minimum cluster size *m*_*clSize*_ to 250 and the local density smoothing factor *m*_*pts*_ to 350. In this framework, points are left unassigned if they reside in sparse regions failing the local density threshold defined by *m*_*pts*_ or if they form clusters smaller than *m*_*clSize*_. To determine a minimum entropy threshold, we identified the cluster with the highest mean entropies and set its minimum values as a cutoff (**Supplementary Fig. 6B**). Only crops with both channel entropies above these thresholds were retained. Using this criterion, we preserved 85.85% of the dataset, corresponding to 19,417 crops (**Supplementary Fig. 6C**). From these, we further excluded crops lacking adjacent neighbours, as they are incompatible with our SimCLR variant, resulting in a final dataset of 19,326 crops (**Supplementary Table S2**). Crops were then resized to 112*112 pixels prior to training. This resolution was chosen to accommodate computational constraints while enabling the use of larger batch sizes than would be possible with the standard 224*224 resolution, which is known to benefit contrastive learning methods such as SimCLR^52^.

### SimCLR-based contrastive learning methods

To systematically characterise cellular phenotypes from fluorescence images without relying on segmentation, we trained contrastive models relying on the SimCLR framework^52^. In this setting, positive pairs are generated by applying independent augmentations to the same image and attracting their representations to one another, whereas negative pairs consist of views from different images whose embeddings are pushed apart. Although effective for diverse natural-image datasets such as ImageNet^53^, this strategy is less well suited to biological images, where samples often exhibit high visual similarity, increasing the risk of false negatives and limiting representation quality^54,68^. To address this limitation and given that positive-pair design critically influences representation quality, we modified SimCLR to include spatially neighbouring crops as additional positive pairs^17,45,46^ (**Fig. 1C**). This strategy actively reduces the information shared by images in the same positive pair, while maintaining shared contextual information, such as local cellular organisation. In this way, the model is encouraged to learn more context-aware and biologically informative features, discarding nuisance variables^16–18,45,56,57^. We refer to this modified framework as SI-SimCLR (Spatially informed SimCLR).

To evaluate this approach, we compared crop-level embeddings, i.e. feature vectors extracted from individual image crops, produced by models trained with different SSL strategies (**Fig. 1D**). We trained two ResNet18 models^69^, of 11 million parameters and embedding dimension 512, from scratch on our astrocyte imaging dataset using either SI-SimCLR or the standard SimCLR algorithm, referred to as *ResNet18 SI-SimCLR* and *ResNet18 standard SimCLR*, respectively. A detailed list of the image transformations used in both SimCLR frameworks, associated hyperparameters, and their application probabilities is provided in **Table S5**. For both frameworks, we trained models for 200 epochs with a batch size of 64 positive pairs (128 individual crops), optimising the NT-Xent contrastive loss^70^, using Adam optimizer^71^ with an initial learning rate of 0.001. This configuration follows established principles of contrastive learning, which favour prolonged training and large batch sizes^52,72^, while adapting reported settings from natural image datasets (1 to 2 orders of magnitude larger) to the scale of our microscopy dataset. Importantly, we validated temperature values t in (0.001, 0.005, 0.01, 0.02, 0.05, 0.1, 0.2, 0.4, 0.6, 0.8, 1.0, 2.0, 3.0), which can critically impact performance. For both models, we selected t=0.1 as this value corresponded to the highest separation of biological classes in the embedding space quantified by the silhouette score per experimental batch (**Supplementary Fig. 1D**). Each model was trained on a single NVIDIA H100 GPU for about 8 hours.

### Additional representation learning baselines

To complete the benchmark of *ResNet18 SI-SimCLR*, we considered four additional baselines detailed below, including the 3 embedding-based methods: *ResNet18 pre-trained, ViT-S uniDINO, ViT-S CellPaint*, and a last method relying solely on handcrafted features (**Fig. 2A, Table S4**).

#### ResNet18 Pre-trained

We used a ResNet18^69^ pre-trained on ImageNet^53^ in a supervised manner to compute baseline embeddings. The ImageNet dataset comprises over one million natural images across 1,000 classes, providing substantially greater visual diversity than our microscopy dataset. The classification head was removed, and features were extracted from the penultimate layer. Input images were resized and normalised using standard ImageNet preprocessing.

#### ViT-S uniDINO and ViT-S CellPaint

We extracted embeddings, without fine-tuning, from two publicly available small Vision Transformer (ViT-S) models, pretrained on CellPainting^73,74^ datasets of multi-cell fluorescence microscopy images with 5 channels. Both models were pretrained using the SSL DINO^75^ framework, but differ in how they process multi-channel information. *ViT-S uniDINO*^*14*^ learns representations independently for each fluorescence channel, which are subsequently concatenated, yielding 768-dimensional embeddings for our two-channel data. Whereas *ViT-S CellPaint*^*14,76*^ processes all channels jointly and outputs 384-dimensional embeddings (**Supplementary Table S6**).

#### Hand-crafted features extraction

We assembled a collection of features to capture texture, intensity, and structural properties of image patches and serve as a baseline against learning-based representations. Similar strategies have proven effective for microscopy image analysis in previous studies^77,78^. In total, 188 features were extracted utilising the scikit-image and scikit-learn Python libraries. The feature set includes Gabor features^79^, Haralick texture features^80^, Tamura features^81^, image pixel intensity distribution (including mean and variance above Otsu’s threshold^82^), spatial moments^83^, Chebychev-based features^84^, edge features based on the Prewitt filter^85^, and Shannon entropy of each channel^86^ (**Supplementary Table S4**). To ensure comparability across scales, all features were z-normalised. Channel-specific features were computed on each of the two image channels.

### Batch effect and biological signal quantification via clustering analysis

To quantify the information captured by different embeddings, we computed the Adjusted Rand Index (ARI) between identified clusters and reference labels. ARI was used to assess separately batch effects (experimental batch labels) and biological signal (ALS/control and TIC treatment), with higher values indicating stronger concordance between cluster assignments and ground-truth categories. Embeddings were first projected in 2D using Uniform Manifold Approximation and Projection (UMAP) with varying hyperparameters (min_dist ∈ {0.001, 0.1}; n_neighbors ∈ {15, 50, 100, 150, 200, 300, 600}) and a fixed initialisation, following established benchmarks^87,88^. KMeans clustering was then applied to the resulting embeddings, using k = 4 as the number of batches when evaluating batch effects (**Fig. 2B**), and k = 4 or 6 when evaluating biological signal within each batch, depending on the number of biological conditions present (**Fig. 2C**). For each model and batch, the maximum ARI obtained across the UMAP parameter sweep was reported.

### Leave-one-batch-out classification

To assess whether ALS and TIC-treated signals generalise across experiments, we trained logistic regressions using a leave-one-batch-out scheme: for each experimental batch, the classifier was trained on all other batches and tested exclusively on the excluded batch (**Supplementary Figs. 2A-B**). All classification tasks were binary: ALS classification was performed using untreated samples as controls, and TIC treatment classification was performed using control samples. Datasets were randomly subsampled to ensure balanced training and test sets across batches and labels, and classifiers were trained using the LBFGS optimizer^89^. Results were averaged across 10 random seeds, controlling subsampling and train–test splits.

### Detection of shared features between ALS and TIC treated astrocytes

To investigate whether ALS and TIC-treated astrocytes share biologically relevant characteristics, we trained an MLP classifier to predict whether a given representation corresponds to TIC treatment and used it to estimate the probability that ALS representations would be classified as TIC-treated (**Figs. 5F,G; Supplementary Fig. 5**). Specifically, for each batch, the dataset was divided into equal, non-overlapping subsets: two subsets consisting of untreated controls, one subset containing untreated ALS samples, one subset containing TIC low-treated controls (if present), and one subset containing TIC high-treated controls. For training, one of the untreated control subsets was combined with one of the other groups (ALS, TIC low, or TIC high). Testing was then performed on the other, non-overlapping control subset combined with the remaining group of interest (either TIC if ALS was used for training, or ALS if TIC was used for training). This approach allowed us to estimate the probability that ALS samples exhibit features similar to TIC-treated cells, and vice versa. For batch E1, each subset contained 508 samples. The MLP classifier comprised a single hidden layer with 100 neurons and was trained for up to 1000 iterations, or until the training loss failed to improve by at least 10^−4^ for 10 consecutive epochs. A batch size of 200 was used. Model selection for testing was performed using five-fold cross-validation.

### Clustering of astrocyte images

To define astrocyte substates in an unsupervised manner, we clustered SI-SimCLR representations. First, we applied UMAP^59^ projections with numbers of nearest neighbours (NN) varying from 15 to 600, similarly than for the previous clustering analyses, but fixing min_dist=0.1 to reduce computational cost as we observed minimal impact on performance. Then we evaluated two clustering strategies, KMeans and Shared Nearest Neighbours (SNN)^90^, on each produced 2D embeddings. The popular KMeans algorithm partitions data into a predefined number of compact, spherical clusters by minimising within-cluster variance, making it computationally efficient but limited in its ability to capture complex structures. In contrast, SNN fits better to our biological embeddings where local relationships and continuous state transitions are important as it enables the robust identification of clusters of varying shape and density by: (i) identifying the k-NN of each point; (ii) computing pairwise similarities based on the proportion of shared neighbours above a threshold t; (iii) performing clustering via DBSCAN, with induced hyperparameters, that automatically detects the number of clusters as densely connected regions. The number of neighbours k controls the scale of local structure, while t sets the stringency of connectivity, jointly determining cluster granularity and robustness to noise. Points in low-density regions that fail to meet these criteria are classified as noise and remain unassigned. We validated k in (50, 75, 100, 125, 150, 175, 200) and t in (0.6, 0.7, 0.8, 0.9). We discarded hyperparameter combinations yielding more than 15% unassigned points. For retained configurations, we computed the missing cluster assignments a posteriori via k-NN majority voting (k=25). Notably, within-cluster variance in handcrafted feature values was low, reflecting the internal homogeneity of the identified substates (**Supplementary Fig. 8**). All clustering results were evaluated using the ARI relative to biological labels and the silhouette score. The best compromise between both metrics was achieved by SNN clustering (k=125, t=0.9) applied to UMAP computed with 15 nearest neighbours, which identified 12 clusters (**Supplementary Fig. 9**).

### Robustness of astrocyte image clusters

For the optimal clustering framework identified above, we assessed robustness to stochastic variability using a consensus clustering analysis. As SNN steps are deterministic, variability was introduced by modifying the initialisation of UMAP embeddings, before running the pipeline 100 times. Global consistency was quantified by the mean pairwise ARI across runs (0.78±0.06), indicating strong reproducibility of our cluster assignments beyond chance. To further visualise consensus structure, we calculated a co-association matrix C, where entries *C*_*i,j*_ represent the frequency of where entries samples i and j sharing a cluster:

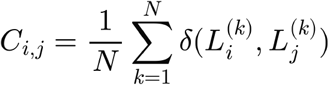

Robust consensus clusters were visualised via a heatmap (**Supplementary Fig. 9B**) by reordering this matrix using agglomerative hierarchical clustering (average linkage) on the dissimilarity 1-C.

### Hierarchical clustering of astrocyte substates

We investigated higher-order relationships between astrocyte sub-states by performing hierarchical clustering on the SNN clusters previously identified (**Fig. 4G**). Specifically, pairwise dissimilarities between clusters were quantified using the Wasserstein distance^91^, which measures the minimal cost of transporting one probability distribution into another. In this context, each cluster was represented as an empirical distribution of samples in the high-dimensional embedding space, with uniform weights assigned to all samples. The Wasserstein distance captures both differences in cluster centroids and discrepancies in their internal geometric structures, providing a distribution-aware measure of inter-cluster dissimilarity. The resulting symmetric distance matrix was then used to construct a hierarchical dendrogram using Ward’s minimum variance^92^.

### Enrichment analysis of astrocyte substates

To evaluate the statistical enrichment of biological classes within the identified clusters, we employed adjusted residuals based on a contingency table framework. First, we constructed a two-way contingency table cross-tabulating cluster assignments and biological classes. Under the null hypothesis of independence between cluster and mutation, we computed expected frequencies using a Chi-squared test^93^. For each cell in the contingency table, adjusted Pearson residuals were calculated to identify specific enrichments while accounting for varying marginal totals. The adjusted residual for row i and column j was computed using the following formula:

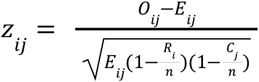

where *O*_*ij*_ is the observed frequency, *E*_*ij*_ is the expected frequency, *R*_*i*_ is the total for row i, *C*_*j*_ is the total for column j, and n is the total number of observations. To control the family-wise error rate across the entire contingency table, a Bonferroni correction was applied for the 72 independent combinations tested. A biological class was considered significantly enriched within a specific cluster if the corresponding adjusted residual was positive and exceeded the Bonferroni-corrected critical value, corresponding to a final threshold of *P*<0.05 (**Supplementary Fig. 3B**).

### Construction of the morphological transition graph

We model the landscape of astrocyte morphological states as a directed probabilistic graph, in which nodes represent phenotypic clusters and edges encode transition probabilities between states (**Fig. 3E**). This framework is designed to capture both the preferred directions of morphological change and the intrinsic ease with which such transitions occur.

Graph construction proceeds in three steps. First, clusters are represented as empirical distributions in the embedding space, enabling a principled geometric formulation within the optimal transport framework. Second, inter-cluster relationships are quantified using partial optimal transport, which identifies sparse and geometry-aware mappings between states. Third, transport-derived quantities are transformed into a stochastic transition matrix using a scale-invariant formulation, thereby yielding a probabilistic description of morphological change dynamics.

#### Clusters as empirical distributions

Let {*c*_1_, …, *c*_12_} denote morphological clusters in the learned embedding space. In order to leverage OT, we model each cluster as an empirical probability measure

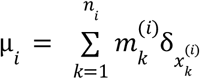

where *n*_*i*_ is the number of samples in the cluster *i*, and 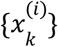 are the corresponding samples, all assigned to a uniform probability mass 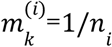. That way, all clusters are assigned a total mass of 1, modelling the a priori assumption that each state is equiprobable, independent of the number of observed samples it contains, since the currently estimated cluster proportions with few samples may be overly influenced by irrelevant technical biases.

#### Estimation of inter-cluster transport

We develop a methodology that enables the estimation of transitions across clusters while accounting for the concurrent nature of morphological variation. This method first assumes that each cluster *c*_*i*_ can independently be a source state and studies how all its biological samples may change, knowing the morphological variations needed to transition to any other possible state. This is modelled by computing the partial Optimal Transport^94^ to move from the cluster probability measure μ_*i*_ to the union of all remaining cluster distributions, denoted as 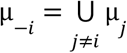. In full generality, partial OT enables comparing distributions with arbitrary masses while enforcing that a total amount of mass 0 ≤ *s* ≤ *min*(|μ_*i*_ |, |μ_−*i*_|) must be transported, by solving the following optimisation problem:

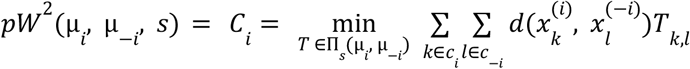

where 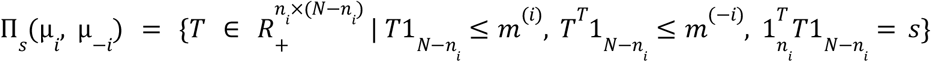 is the set of admissible transport matrices with total transported mass *s* and marginals bounded by mass vectors *m*_^(*i*)^_ and *m*_^(−*i*)^_. Here to ensure that all the mass in μ_*i*_ is transported, we fix *s* = 1, so that the constraint on *s* is saturated and necessarily *T* satisfies 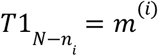 *and* 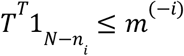. Importantly, this process enables the selection of a sparse set of points in μ_−*i*_, with a total mass of 1 instead of 11. Therefore, denoting *T*^(*i*)^ the partial OT matrix, we consider as first estimations of transition probability to leave *c*_*i*_, the cumulated mass arriving to the cluster *c*_*j*_:

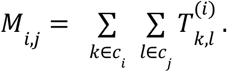

Conceptually, this partial OT matching estimates the minimal averaged morphological variation (measured by Euclidean distances) such that each point in μ_*i*_ is transformed into at least one point in μ_−*i*_. Importantly, we stress that such a selection model accounts for the dynamic nature of morphological variation, which cannot be easily modelled using classical balanced OT via the Wasserstein distance. In particular, it would require normalising the mass of μ_−*i*_ to the one of μ_*i*_ = 1, such that every point in μ_−*i*_ would receive mass, which would come down to admitting larger magnitudes of morphological variation and being exposed to a higher sensitivity to noise. Alternatively, one could envision Wasserstein matchings between all pairs (μ_*i*_, μ_*j*_) that would amount to assuming the existence of a direct “straight line” (formally, a geodesic) between both morphological states, without accounting for the fact that points must pass through other observed morphological states, thereby defying the Waddington paradigm.

#### Scale-invariant formulation of transition propensity

For a given state *c*_*i*_, the cumulated transported masses *M*_*i,j*_ provide how points are supposed to move to the other states, while enforcing all points to move, i.e. considering *c*_*i*_ as a source. However, across all real potential sources, as 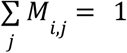, the transported masses do not express how difficult it is to move points from one state to another across source states. This notion is captured by the partial OT costs, such that the smaller *C*_*i*_ is, the more likely it is to move from state *i*. We then aim to define an escape probability from a state *i* based on these transportation costs. Importantly, *C*_*i*_ depends on the geometric structure of the embedding space and therefore its absolute scale is not intrinsically meaningful, and only relative differences across clusters should be interpreted. Therefore, we consider scaling costs linearly via a min-max scaling. However, this methodology would lead to unrealistic extreme escape probabilities for the states achieving either *C*_*max*_ = *max*_*i*_ *C*_*i*_ or *C*_*min*_ = *min*_*i*_ *C*_*i*_. Therefore, we first bound the transport costs between their 10% and 90% quantiles, denoted *q*_10_ and *q*_90_, such that the escape probability from state *i* reads as

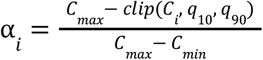

This formulation has three key properties: it is strictly in ]0, 1[ for all states, monotonic in the transport cost and invariant to the scale of *C*_*i*_. Biologically, *α*_*i*_ can be interpreted as the propensity of a morphological state to undergo transition under perturbation. Consequently, the final row-stochastic transition matrix *P* between states is defined as:

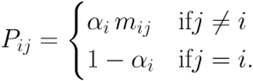

so that it defines a valid Markov process over morphological states. Notice that the inclusion of self-loops reflects the biological reality that most cells do not change state at every observation step, especially in stable or low-plasticity conditions.

### Pruning of the morphological transition graph for visualization

To ease visualisation, we focused on high-frequency signals by pruning the edges below the third quartile *Q*_3_, following the statistical thresholding approach of PAGA^95^, retaining by construction the top 25% of transitions and yielding a sparse, biologically interpretable graph. States whose entire row is zeroed by pruning are treated as terminal absorbing states with no significant outgoing flux.

### Graph-theoretic metrics

We characterised the morphology transition graph using a set of complementary state-level metrics designed to capture directional flux, transition diversity, and topological roles within the phenotypic landscape. All metrics were computed on the off-diagonal transition probabilities *P*_*i,j*_ for *i* ≠ *j*, thereby isolating inter-state dynamics from state persistence encoded by self-loops.

#### Net flow

We measure the overall directionality of transitions from a state *i* via its net flow^96^, defined as the difference of outgoing and incoming probability masses 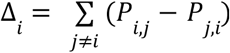. Positive values (Δ_*i*_ > 0) indicate that a state exports more transition probability mass than it receives (progenitor-like behaviour), whereas negative values (Δ_*i*_ < 0) indicate that it preferentially accumulates incoming transitions (attractor-like behaviour).

#### Generalised entropy of unnormalised transitions

To quantify transition diversity without enforcing probability normalisation, we define a generalised entropy directly on the unnormalised outgoing transition weights *P*_*i,j*_ for *i* ≠ *j* (excluding self-loops). We compare this empirical distribution to a uniform reference measure over the *K* − 1 = 11 possible target states. Then we define a generalised entropy based on the generalised Kullback–Leibler divergence between both distributions, accounting for their difference in total mass: 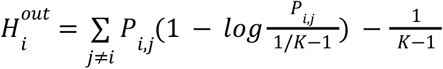. This quantity measures how far the observed unnormalised transition profile of a morphological substate deviates from a uniform reference over all possible target states. Low values indicate concentrated or selective transitions (low diversity in morphological changes), whereas high values indicate broadly distributed transition mass (high diversity). Importantly, this formulation does not require stochastic normalisation and therefore preserves information about absolute transition strength, such that substates remain comparable. A similar quantity denoted 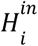 is derived to measure the diversity of incoming transition weights.

#### Betweenness centrality

Betweenness centrality *B*_*i*_ quantifies the extent to which a node lies on shortest paths between other states in the graph^97^. Shortest paths were computed using edge costs defined as the inverse of the transition probability, so that high-probability transitions correspond to short distances. States with high betweenness centrality act as critical intermediates connecting different regions of the morphological landscape. These states may function as transitional hubs through which multiple phenotypic trajectories pass. Unlike net flow or entropy, betweenness captures global topological structure rather than local transition patterns.

### Primary state classification

Morphological substates were classified into three primary functional archetypes using a hierarchical decision rule based on betweenness centrality and net flow (**Fig. 3F**). The key insight underlying this scheme is that betweenness centrality and net flow capture complementary and partially independent aspects of a node’s role in the transition graph. Betweenness is a topological property: a node with high betweenness lies on many shortest transition paths between other substates, which by definition requires it to both receive and redistribute flux. A genuine source or sink, by contrast, participates predominantly in one-directional flux and therefore does not serve as a topological intermediary. Net flow is a directional property: it measures whether a node is a net exporter or importer of transition mass, independently of its topological centrality. The classification proceeds as follows. Let *B* = *median*({*B*_*i*_}) denotes the median betweenness across all states. A state is classified as:

#### Transient

If *B*_*i*_ ≥ *B*, indicating that the node lies on a disproportionate number of shortest transition paths and therefore functions as a topological crossroads between other substates. Note that a node with high net flow and high betweenness is classified as Transient rather than Source, because high betweenness implies the node both receives from and sends to diverse substates — consistent with a transitional rather than a purely generative role.

#### Source/Progenitor

if *B*_*i*_ < *B* and Δ_*i*_ > 0, indicating a net sender that is not a topological intermediary. Such nodes export more flux than they receive without serving as bottlenecks on transition paths.

#### Sink/Attractor

if *B*_*i*_ < *B* and Δ_*i*_ < 0, indicating a net receiver that is not a topological intermediary. Such nodes accumulate incoming flux without substantially redistributing it to other substates.

### Complementary state classification

Two additional binary flags were assigned orthogonally to the primary classification, such that any substate can carry either, both, or neither flag independently of its primary archetype.

*Hub*. A substate is flagged as a hub if its betweenness *B*(*i*) exceeds the first quartile of the betweenness distribution and it maintains at least three non-zero connections in both the incoming and outgoing directions. The degree conditions ensure that hub status reflects genuine bidirectional connectivity rather than high betweenness arising from a sparse graph with few alternative paths.

#### High-plasticity states

A substate is flagged as high-plasticity if its outgoing transition entropy 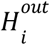 exceeds the median of the corresponding distribution and it has at least three non-zero outgoing connections. A high-plasticity state is one that disperses outgoing flux broadly across many downstream substates, capturing the biological notion of phenotypic plasticity, i.e. the capacity to give rise to a diversity of downstream morphological configurations. Hub states are a subset of high-plasticity states in many cases, but the two flags are not equivalent: a hub requires both incoming and outgoing diversity, whereas a high-plasticity state requires only outgoing diversity and may have low incoming connectivity.

### Transition between control and ALS astrocytes

To characterise transitions from control to ALS astrocytes, we applied partial OT described in section “*Construction of the morphological transition graph*” to control (all conditions) and ALS (basal) high-dimensional embeddings, seen as source and target distributions, respectively (**Figs. 5D-E)**. For our problem, we defined the transported mass to match the smaller distribution between control (all conditions) and ALS (basal). This ensured that every point in the smaller sample was accounted for and mapped to a corresponding subset in the larger population. The resulting sparse point-to-point transport plan was subsequently aggregated by summing the transport mass flowing between all embeddings of a specific source cluster and those of a target cluster. This produced a coarse-grained transition matrix, which was column-normalized, such that the total incoming flow to each target cluster summed to 1. This normalisation allowed us to quantify the proportional contribution of each TIC condition to each ALS state.

### Molecular analysis

#### Gene expression preprocessing and filtering

Whole-transcriptome quantification was performed using Kallisto^98^, and transcript-level estimates were aggregated to the gene level using the GENCODE v44 annotation. Genes with zero counts across all samples were removed. To distinguish reliably expressed genes from background noise, a two-component Gaussian Mixture Model (GMM) was fitted to log_2_-transformed expression values for each sample. The first component (“background”) represents lowly expressed genes corresponding to noise, whereas the second component (“foreground”) represents reliably expressed genes. An expression threshold was defined as the 95th percentile of the background distribution. Genes were retained if their expression exceeded the sample-specific threshold in at least one sample. Following filtering, the gene expression matrix was quantile-normalised across samples to account for differences in library size and composition.

#### Hierarchical clustering and dimensionality reduction

Hierarchical clustering was performed on the filtered and normalised gene expression matrix using complete linkage and Spearman correlation as the distance metric (1 − ρ). Principal Component Analysis (PCA) was performed on the expression matrix using a Singular Value Decomposition (SVD) approach, and the first component was removed by subtracting its contribution from the original data.

#### Pathway activity estimation

For each patient and each Gene Ontology Biological Process (BP) term, the average expression of genes annotated in the BP was calculated. To account for pathway size effects, gene sets of identical size were randomly sampled from genes not belonging to the BP to generate a background distribution of mean expression values. This bootstrap sampling was repeated 100 times for each BP. The observed BP mean expression was then converted to a z-score relative to the background distribution, from which a p-value was derived. Finally, p-values were transformed using a −log10 scale and assigned a sign based on pathway regulation – positive for enrichment and negative for depletion – resulting in a matrix of pathway activity scores for each GO BP term across all samples. Finally, the pathway activity scores were averaged across samples within each group (treatment and mutation).

#### Identification of condition-specific pathways

PCA was performed on the pathway activity matrix using a SVD approach, and the first component was removed by subtracting its contribution from the original data to reduce global effects. The resulting distribution of percentage of variance explained per component is shown in **Supplementary Fig. 3F**. To identify components associated with treatment status (Untreated, TIC-low, TIC-high), linear mixed-effects models were fitted with treatment as a fixed effect and mutation as a random effect. The first principal component (PC1), explaining 37.1% of the variance, showed the strongest association with treatment (**Supplementary Fig. 3G**). Similarly, to identify components associated with mutation status (CTRL vs VCP), each left singular vector was evaluated using a linear mixed-effects model with mutation as a fixed effect and treatment as a random effect. The third principal component (PC3), explaining 16.8% of the variance, best captured mutation-associated variation (**Supplementary Fig. 3H**). For both the first (PC1) and third (PC3) principal components, the 50 pathways with the highest positive and 50 with the most negative contributions (loadings) were selected for further analysis.

#### Visualising pathway relationships

For each component (PC1 and PC3), the resulting 100 pathways were manually grouped into broader functional categories, as described in **Table S7**. To visualise relationships among pathways within the same category, a pathway network was constructed in which each node represents a GO term. Node size was proportional to the number of genes annotated to the pathway. Edges between nodes were defined based on gene overlap between pathways, quantified using the Jaccard similarity index. For each pair of pathways, the Jaccard index was calculated as the size of the intersection of their gene sets divided by the size of their union. An edge was drawn whenever the Jaccard similarity exceeded 0.01, indicating shared gene membership. Edge width was proportional to the Jaccard similarity score, with thicker edges representing greater gene overlap between pathways. The network layout was generated using a circular layout algorithm that positioned nodes evenly around a circle to provide a clear and systematic visualisation of pathway relationships.

## Data availability

The raw and processed images and respective annotations have been deposited on Zenodo under the accession number http://doi.org/10.5281/zenodo.19553134. All sequence data for this project has been deposited at NCBI GEO database under accession number GSE331442.

## Code availability

The source code is available on Github at: https://github.com/AI-for-RNA-Biology/SI-SimCLR

## AUTHOR CONTRIBUTIONS

Conceptualization, R.L., R.P., C.V.C.; Formal Analysis, E.M., L.F. D.T. performed the experiments. Investigation, E.M., G.E.T., J.N., H.C., P.K., O.Z. D.M.T.; Writing – Original Draft, R.L., R.P., E.M., C.V.C.; Writing – Review & Editing: all authors; Resources, R.L., R.P., P.F., V.U..; Visualization, E.M., L.F..; Supervision, R.L,, R.P., C.V.C.

## ACKNOWLEDGMENTS

The authors wish to thank the patients for fibroblast donations. We are grateful for the help and support provided by the team of developers at Idiap.

## Legends of Supplementary Tables

Supplementary tables are accessible here (public link) https://docs.google.com/spreadsheets/d/1qK7goOPVLzzaRhWfnNdC7-EffAX2_T-ajpt27-z3tPk/edit?usp=sharing

**Table S1** | Dataset statistics before pre-processing (z-stack and channels merging only). Details regarding specific treatments and cell lines are provided for each experimental batch. Cell lines are designated as either CTRL (control) or MUT (VCP).

The “N. of images” column indicates the total number of 1080×1080 RGB images generated via maximum intensity projection of the z-stacks.

**Table S2** | Number of crops per class (mutation and treatment). Number of corresponding wells and 270×270 image crops for each experimental batch, mutation, and treatment, after excluding low-content crops based on entropy thresholding.

**Table S3** | RNA sequencing data. Sample ids with their corresponding library type, cell line, treatment and mutation status.

**Table S4** | Hand-crafted features. Individual hand-crafted features grouped by category.

**Table S5** | **Data augmentations applied to images for SimCLR and SI-SimCLR**.

Each transformation was applied stochastically with a probability specified in the “Probability” column. For rotation, hyperparameter set denotes the options from which a single value is sampled uniformly. Color jitter parameters denote the uniform distributions [min, max] from which scaling factors (for brightness, contrast, and saturation) and additive shifts (for hue) are sampled. Abbreviations: N/A, not applicable. **Table S6** | **Overview of embedding sets and feature extractors**. Architectural features, training algorithms, and training datasets used to generate the different embedding sets. Hand-crafted features do not utilize a learned feature extractor or training data (denoted as N/A). Abbreviations: s.c., single channel; N/A, not applicable.

**Table S7** | **Biological Pathway groups**. Top 100 GO pathways contributing to the first (left) and third (right) principal components of pathway-PCA, manually grouped into broader functional categories.

## Supplementary Figures

**Supplementary Figure 1.**
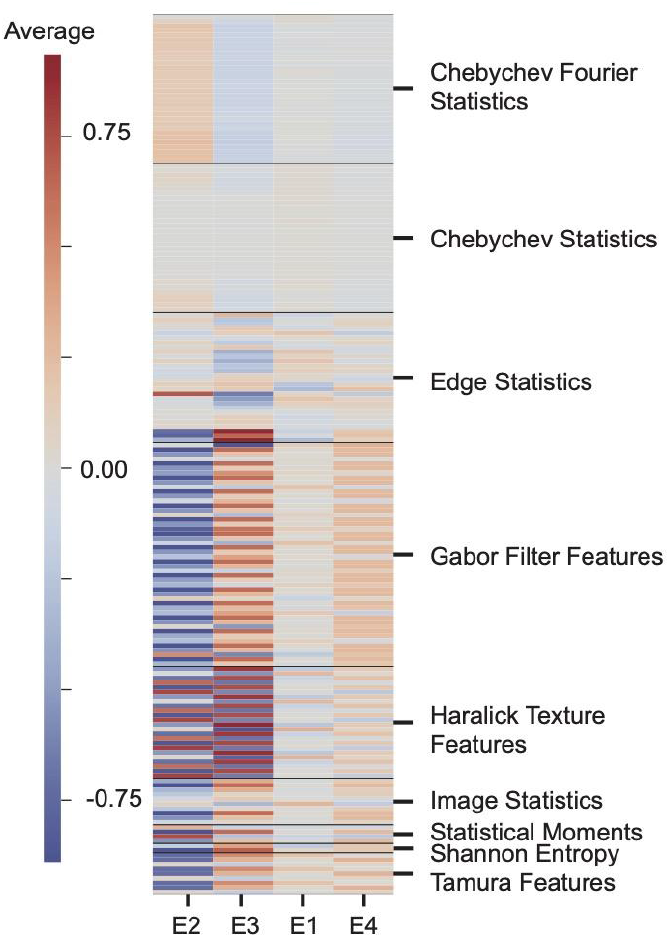
Heatmap displaying the average value of hand-crafted features in each experimental batch. Labels on the right of the heatmap indicate the broad categories in which the features were grouped. The complete list of hand-crafted features is described in Supplementary Table S4.

**Supplementary Figure 2.**
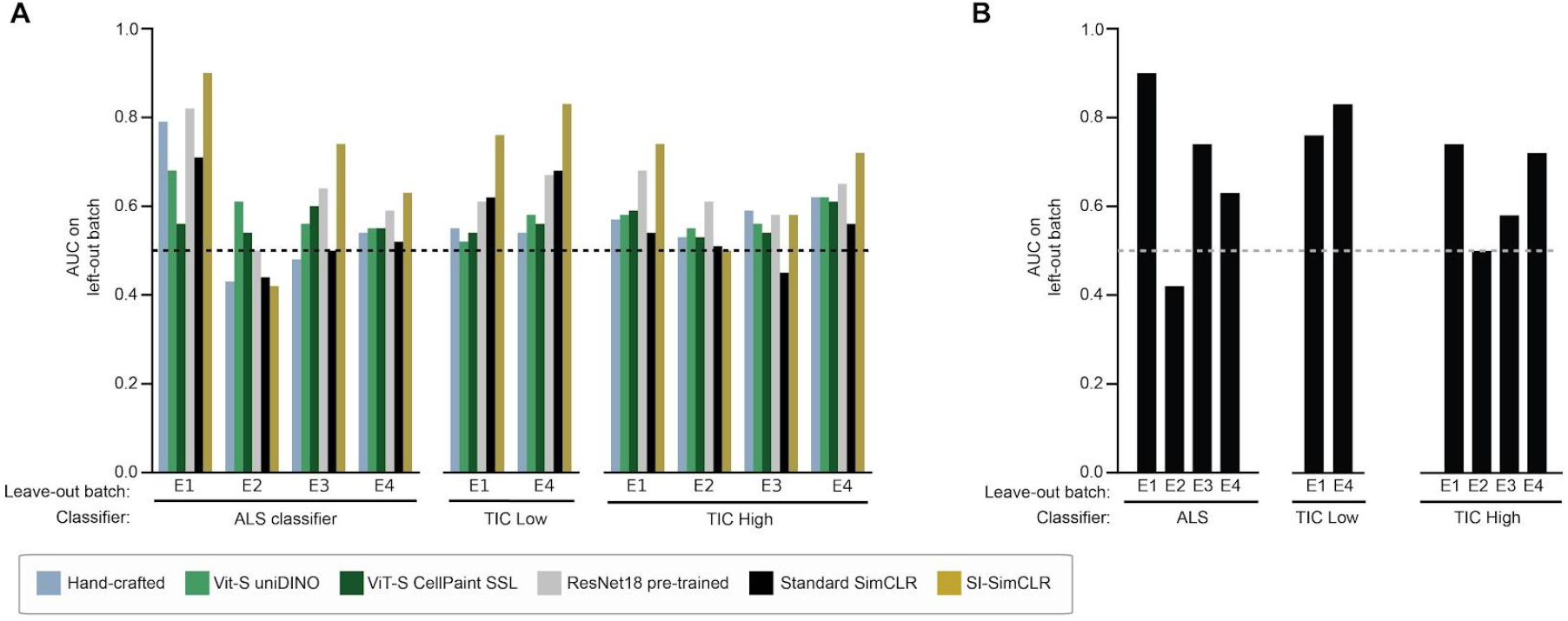
**A**. Barplots showing the generalisation performance (AUC scores) of logistic regression classifiers using a leave-one-batch-out evaluation strategy. Individual bars represent the AUC score achieved on each specific held-out batch by training on the three remaining ones. Models were trained to distinguish: control vs. ALS astrocytes (left), control untreated vs. TIC low (middle), and control untreated vs. TIC high (right). Dashed lines represent AUC=0.5.**B**. Same as **A**, but only for the SI-SimCLR model.

**Supplementary Figure 3.**
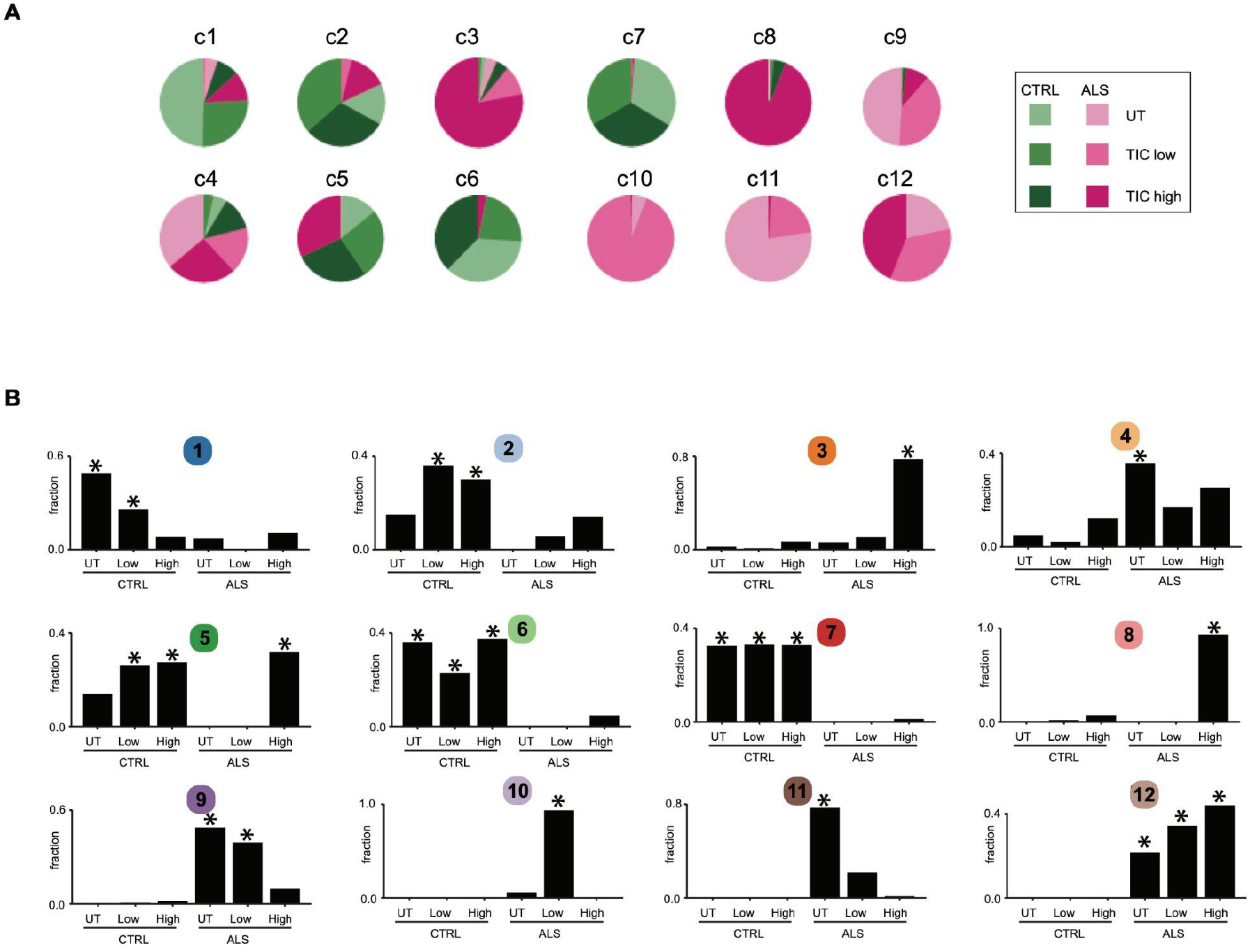
**A**. Pie charts showing the relative abundance of each biological condition for each identified morphological substates. **B**. Barplots showing the relative abundance in each biological condition (genotype and inflammation) for each identified morphological substates. * P<0.05 with Fisher enrichment test.

**Supplementary Figure 4.**
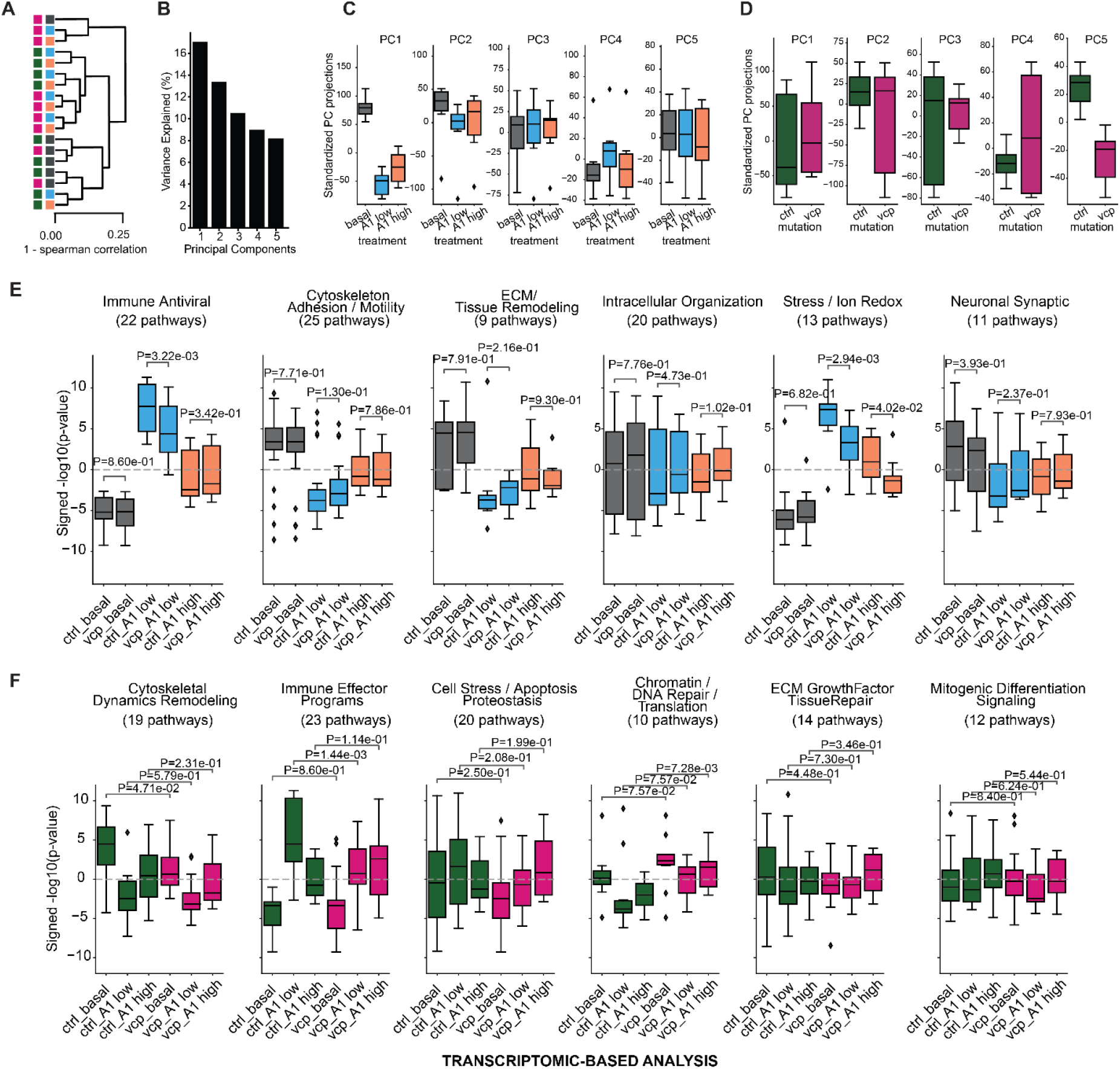
**A**. Hierarchical clustering performed on the filtered and normalised gene expression matrix using complete linkage and Spearman correlation as the distance metric (1 − ρ). Samples are coloured by mutation (left) and treatment (right). **B**. Percentage of variance explained by each principal component (PC) derived from the gene expression matrix using PCA (see *Materials and Methods*). **C**. Box plots of gene expression principal component projections across samples, coloured by treatment. **D**. Same as (G) coloured by mutation. **E**. Pathway groups derived from the top 100 contributing pathways to the pathway activity principal component associated with treatment (see *Materials and Methods*). Boxplots show the distribution of enrichment scores (signed −log_10_(p-values)) for pathways within each group across the six experimental conditions. **F**. Same as I for the principal component associated with mutation.

**Supplementary Figure 5.**
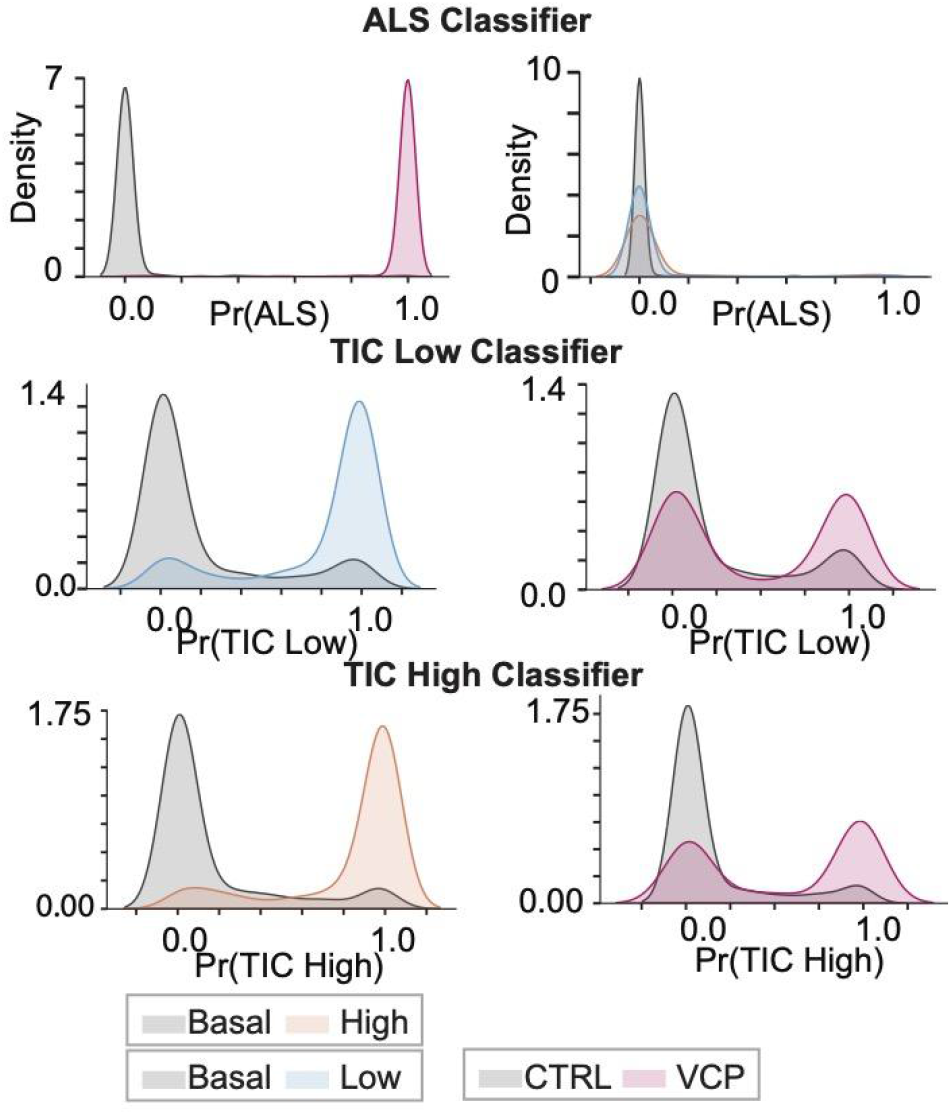
Predicted probability distributions computed by an ALS classifier (left column), a TIC low classifier (middle column) and a TIC high classifier (right column). Top row: MLP were trained via supervised binary classification to distinguish untreated controls from the biological classes specified in the corresponding legends. Bottom row: These models were then applied to unseen, non-overlapping subsets of alternative conditions (bottom row). Specifically, the ALS classifier evaluated TIC samples (bottom of left column), while the TIC classifiers evaluated ALS samples (bottom of middle and right columns).

**Supplementary Figure 6.**
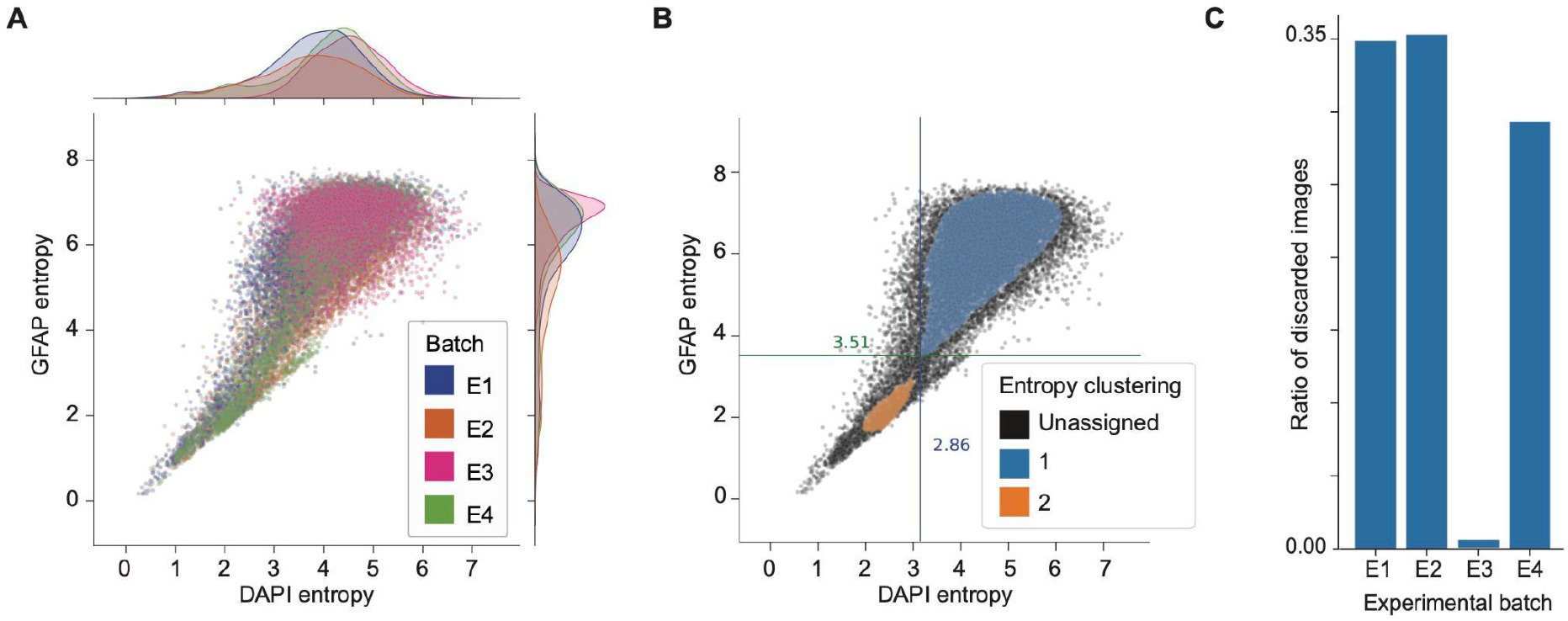
**A**. Scatterplot of Shannon entropy values computed for DAPI (x-axis) and GFAP (y-axis) channels across all images. Marginal distributions are shown along the top and right axes. Points are colored by experimental batch, illustrating batch-wise variability in image information content. **B**. Same as in (A), with points colored according to HDBSCAN^67^ clustering. Thresholds used to identify and discard low-content images are indicated by vertical (DAPI) and horizontal (GFAP) lines. These thresholds were defined based on the lowest entropy values within the cluster exhibiting the highest mean entropy. **C**. Bar plot showing the proportion of images discarded per experimental batch based on the entropy thresholds defined in (B).

**Supplementary Figure 7.**
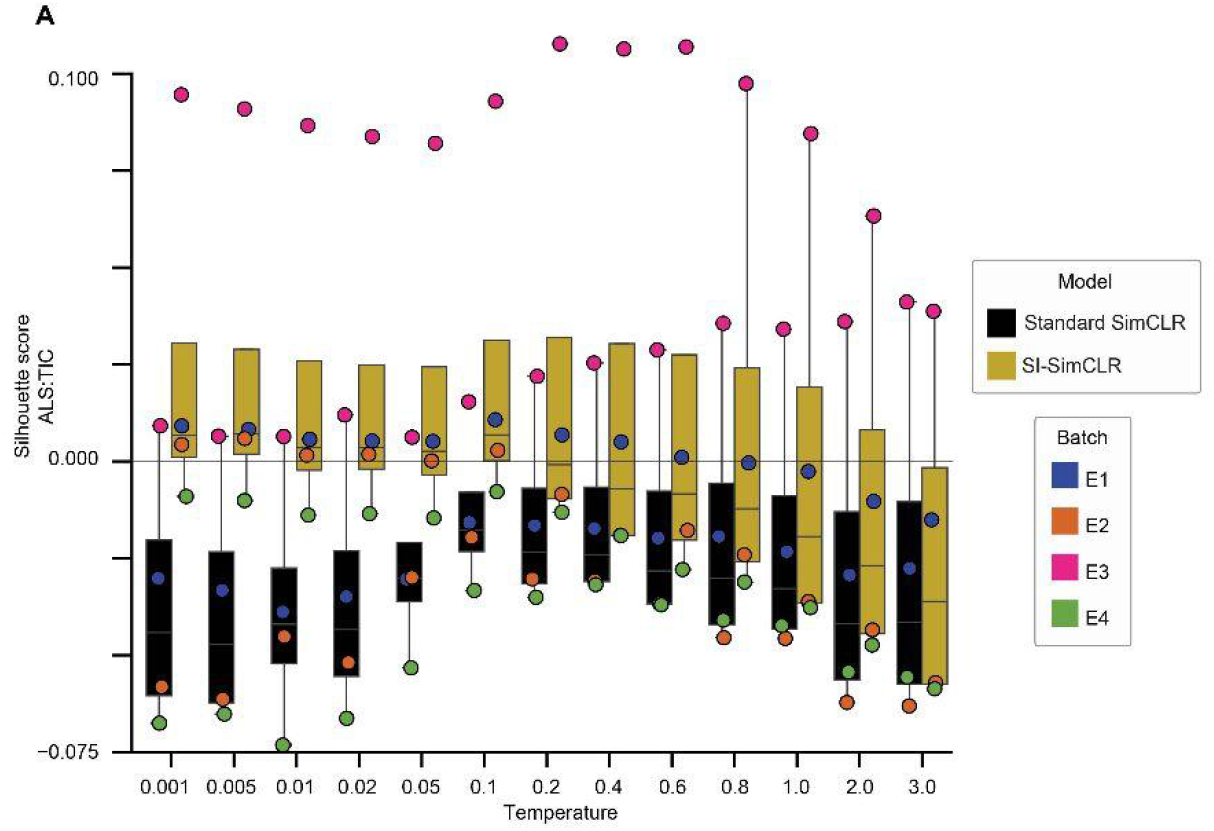
Boxplot of silhouette scores for varying temperatures for SimCLR training. For each training algorithm and temperature value, a ResNet18 was trained on the filtered dataset. Subsequently, we computed the Silhouette score^100^ for each experimental batch using biological conditions (ALS/control, TIC treatment) as labels.

**Supplementary Figure 8.**
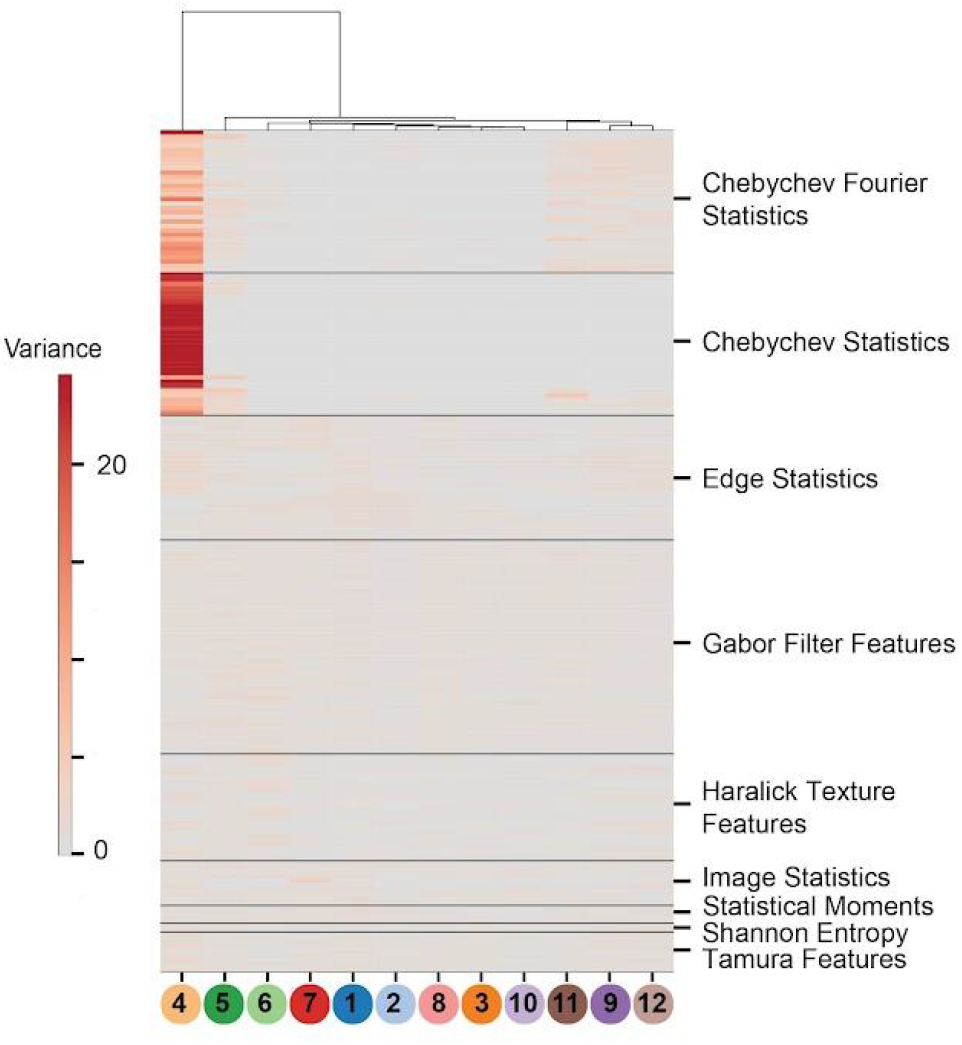
Heatmap representing variance of hand-crafted features in SNN clusters. Labels on the right of the heatmap indicate the broad categories in which the features were grouped. The complete list of hand-crafted features is described in Supplementary Table S4.

**Supplementary Figure 9.**
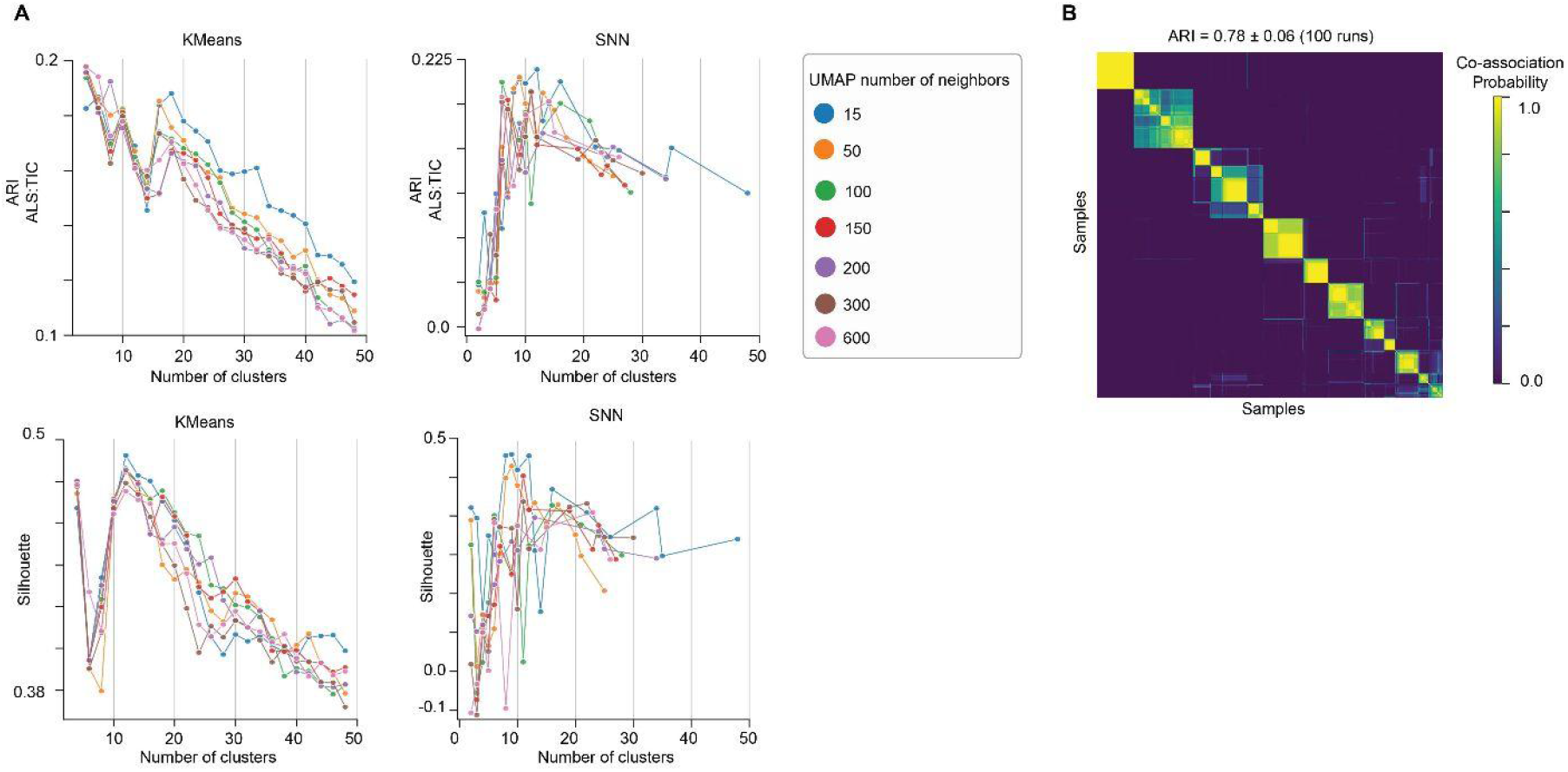
**A**. Evaluation of clustering algorithms. Score values are plotted against the resulting number of clusters. Left column: KMeans^101^ clustering. Right column: Shared Nearest Neighbors (SNN)^90^. Top row: Silhouette score^100^. Bottom row: Adjusted Rand Index (ARI)^60^ comparing cluster labels and known biological classes. Lines are coloured according to the number of neighbours used in the UMAP dimensionality reduction step. Circles indicate the individual clustering runs. **B**. Heatmap representing the co-association matrix of clustering assignments across multiple iterations with different UMAP random seeds. The colour scale indicates pairwise co-association frequency, ranging from 0 (samples were never assigned to the same cluster) to 1 (samples were always assigned to the same cluster). Both axes are sorted according to the final consensus cluster labels derived from hierarchical average-linkage clustering.

